# Plant speciation in the Namib Desert: origin of a widespread derivative species from a narrow endemic

**DOI:** 10.1101/2022.01.04.474907

**Authors:** Joseph J. Milton, Matthias Affenzeller, Richard J. Abbott, Hans P. Comes

## Abstract

**Background:** Parapatric (or ‘budding’) speciation is increasingly recognized as an important phenomenon in plant evolution but its role in extreme (e.g. desert) environments is poorly documented.

**Aims:** To test this speciation model in a hypothesized sister pair, the Southwest–North African disjunct *Senecio flavus* and its putative progenitor, the Namibian Desert endemic *S. englerianus*.

**Methods:** Phylogenetic inferences were combined with niche divergence tests, morphometrics, and experimental-genetic approaches. We also evaluated the potential role of an African Dry-Corridor (ADC) in promoting the hypothesized northward expansion of *S. flavus* (from Namibia), using palaeodistribution models.

**Results:** Belonging to an isolated (potential ‘relict’) clade, the two morphologically distinct species show pronounced niche divergence in Namibia and signs of digenic-epistatic hybrid incompatibility (based on F_2_ pollen fertility). The presence of ‘connate-fluked’ pappus hairs in *S. flavus*, likely increasing dispersal ability, is controlled by a single gene locus.

**Conclusions:** Our results provide support for a rare example of ‘budding’ speciation in which a *wider-*ranged derivative (*S. flavus*) originated at the periphery of a *smaller*-ranged progenitor (*S. englerianus*) in the Namib Desert region. The Southwest–North African disjunction of *S. flavus* could have been established by dispersal across intermediate ADC areas during periods of (Late) Pleistocene aridification.

## Introduction

A long-held view in plant evolution research is that genetic divergence within species is promoted in arid and semi-arid regions due to high evels of population subdivision and isolation (Stebbins 1952, 1972, 1974; Axelrod 1967, 1972). Moreover, cycles of aridity interspersed by periods of more mesic conditions, during climatic oscillations, might be expected to trigger cycles of geographical isolation and renewed contact of species populations, creating conditions which further promote speciation (He et al. 2019). It is not surprising, therefore, that plant speciation and radiations have been shown to be particularly evident in arid regions during the Quaternary when cycles of aridity interspersed by pluvial stages likely occurred (Kadereit and Abbott 2021). Despite this, few studies of plant speciation in arid areas have been undertaken and none as far as we know in sub-Saharan Africa.

Over the last decades, considerable empirical evidence has accumulated to suggest that ‘peripheral isolate’ speciation (*sensu* Mayr 1963) might be common in plants (Anacker and Strauss 2014, and references therein). According to this model, hereafter termed ‘budding’ speciation, a derivative species forms near the edge of a progenitor’s range, leaving the latter almost unchanged (Stebbins 1974; Grant 1981; Gottlieb 2004; Levin 2004; Crawford 2010; Lopéz et al. 2012; Otero et al. 2019, and references therein). Following Anacker and Strauss (2014) a number of predictions arise from this model: (1) young sister species are expected to have parapatric ranges with very different sizes, that is, a *larger*-ranged progenitor and a *smaller-*ranged derivative (but see below); (2) the progenitor would be expected to be initially poly- or paraphyletic with respect to the derivative (e.g. due to incomplete lineage sorting and/or gene flow), while reciprocal monophyly might be attained only with time (e.g. due to lineage sorting, the build-up of reproductive isolation (RI) barriers or extinction); (3) especially for sister species occurring in environments of substantial edaphic and/or climatic heterogeneity, RI mechanisms should encompass ecological isolation due to strong divergent selection across habitats (see also Schluter 2001; Givnish 2010); (4) those ecological differences, in turn, might further promote prezygotic isolation (e.g. shifts in phenology, pollinators, mating system) and/or postzygotic isolation of either an extrinsic type (e.g. due to low adaptation of hybrids to parental habitats; Richards et al. 2016) or an intrinsic kind (e.g. due to reduced hybrid fertility/viability caused by ‘genic’ incompatibilities, Orr and Turelli 2001; Johnson 2008; Satogankas et al. 2020); and (5) ‘budding’ speciation in plants should frequently be accompanied by trait changes that can generate RI more or less instantaneously, such as polyploidization and/or shifts from outcrossing to selfing (Abbott 2017; Slopa et al. 2020).

Studies that have explicitly tested hypotheses of budding speciation in plants remain few in number, with most focused on species in temperate or subtropical ecosystems, while examples from ‘extreme’ environments (e.g. hyper-arid deserts) are rare (e.g. Wang Q et al. 2013; Luebert et al. 2014; Ma et al. 2018; Singhal et al. 2021). Furthermore, little attention has been paid to *smaller-*ranged progenitor and *wider-*ranged derivative species (e.g. Kadereit 1984a; Kadereit et al. 1995), possibly, because of the view that asymmetrical gene flow from central to peripheral populations often retards (or ‘swamps’) local adaptation and expansion of range limits (Mayr 1963; Grant 1981; Kirkpatrick and Barton 1997; Levin 2000; Sexton et al. 2009). Nonetheless, some authors have suggested that, even if limited, such gene flow could provide “the crucial influx of new mutations on which local selection must operate” (c.f. Givnish 2001). Moreover, peripheral populations are often pollen limited due to reduced mate availability and/or pollinator abundance (Busch 2005), which in turn might favour the evolution of selfing and/or dispersal ability (e.g. Baker 1955; Barrett 2014; Iritani and Cheptou 2017).

In the present study, we examine the possibility of budding speciation in a hypothesized sister pair of *Senecio* L. (Asteraceae, Senecioneae; c. 1,200 spp., Walter et al. 2020) involving the Namibian Desert endemic *S. englerianus* O.Hoffm. and its putative derivative species, *S. flavus* (Decne.) Sch.Bip., a (semi-)desert species with disjunct distribution between Southwest and North Africa (Alexander 1979; Kadereit 1984b; Coleman et al. 2003; Figure 1(A)).

**Figure 1.**
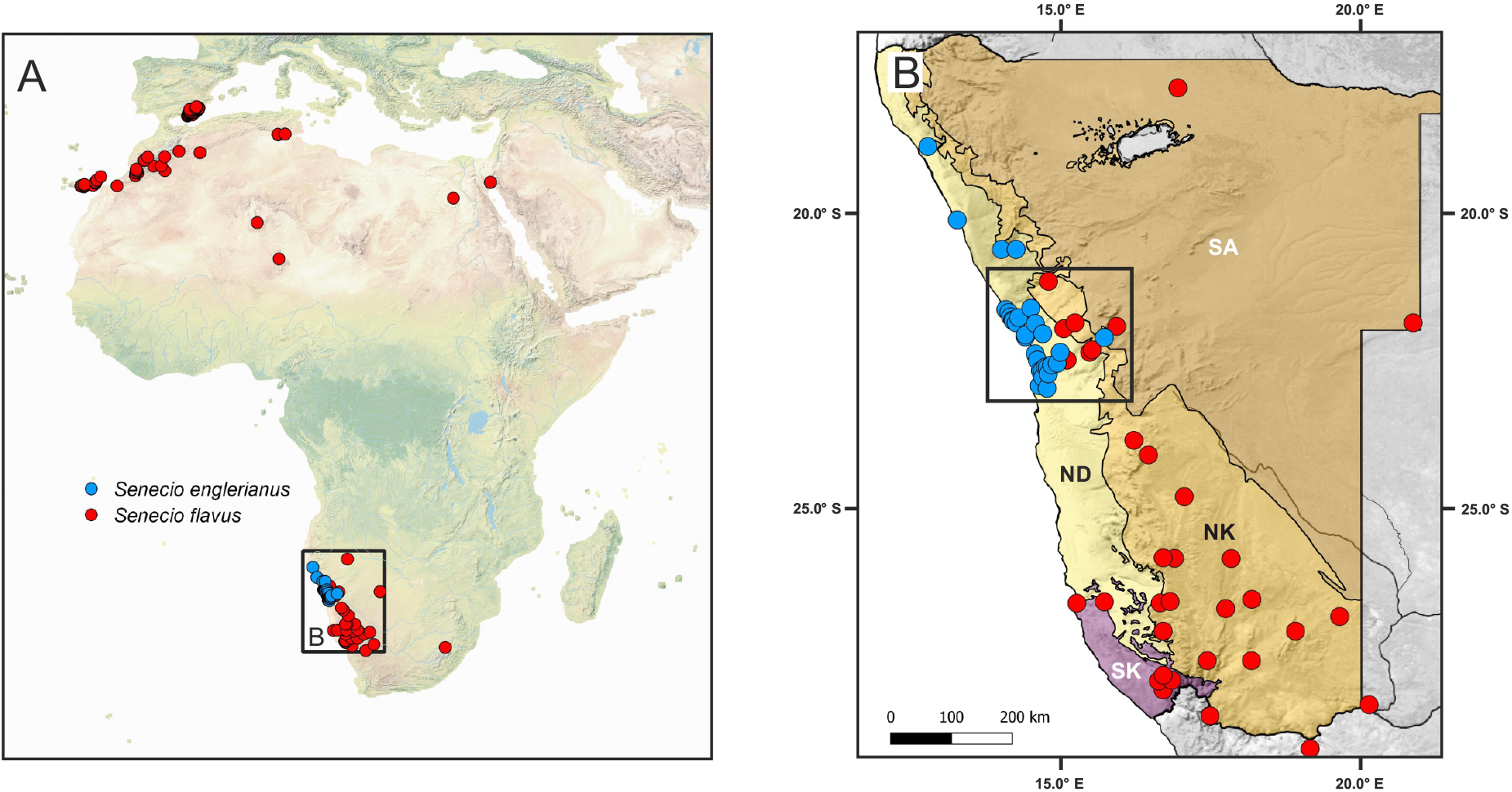
(A) Map of the geographical distributions of the Namibian Desert endemic *Senecio englerianus* (blue dots) and *S. flavus* (red dots) based on geo-referenced locality records (N = 30 and 150, respectively; see text for details). (B) Map showing the species’ distributions across the four major biomes (ecoregions) of Namibia (according to Irish 1994) and adjacent areas (northwestern South Africa: only *S. flavus*). The black retangle indicates the area of species range overlap in the Central Namib Desert and its hinterland (approximately south of Mt. Brandberg to Khan River). Biome abbreviations: ND, Namib Desert; NK, Nama-Karoo; SK, Succulent Karoo; SA, Savanna.

In Namibia, four biomes (or ecoregions) can be distinguished (Irish 1994): Desert, Nama-Karoo, Succulent Karoo, and Savanna (see Figure 1(B)). Based on available locality records (see Materials and Methods), *S. englerianus* is largely restricted to the hyper-arid, fog- nourished coastal plains of the Northern (sandy) and (mostly gravelly) Central Namib Desert (c. 0–870 m a.s.l.), affected by relatively cool temperatures and highly unpredictable rainfall (Giess 1981; Craven and Vorster 2006; Jürgens et al. 2013; Figure 2(A)). By contrast, *S. flavus* is mainly distributed at higher elevations (c. 190–1700 m a.s.l.) in the (semi-)arid Nama-Karoo biome of low-shrub vegetation, which occupies most of the interior plateau region of southern Namibia (extending into northwestern South Africa) but also stretches northwestward in a narrow band along the foot of the ‘Namib Escarpment’ (Figure 2(B)). In addition, *S. flavus* is reported from the Succulent Karoo coastal belt of Southwest Namibia and, very rarely, from the tree and shrub Savanna biome of northern and eastern Namibia (see also Foden et al. 2011). Notably, *S. englerianus* and *S. flavus* occur in para-/sympatry in the interior Central Namib Desert and its hinterland (see black rectangle in Figure 1(B)).

**Figure 2.**
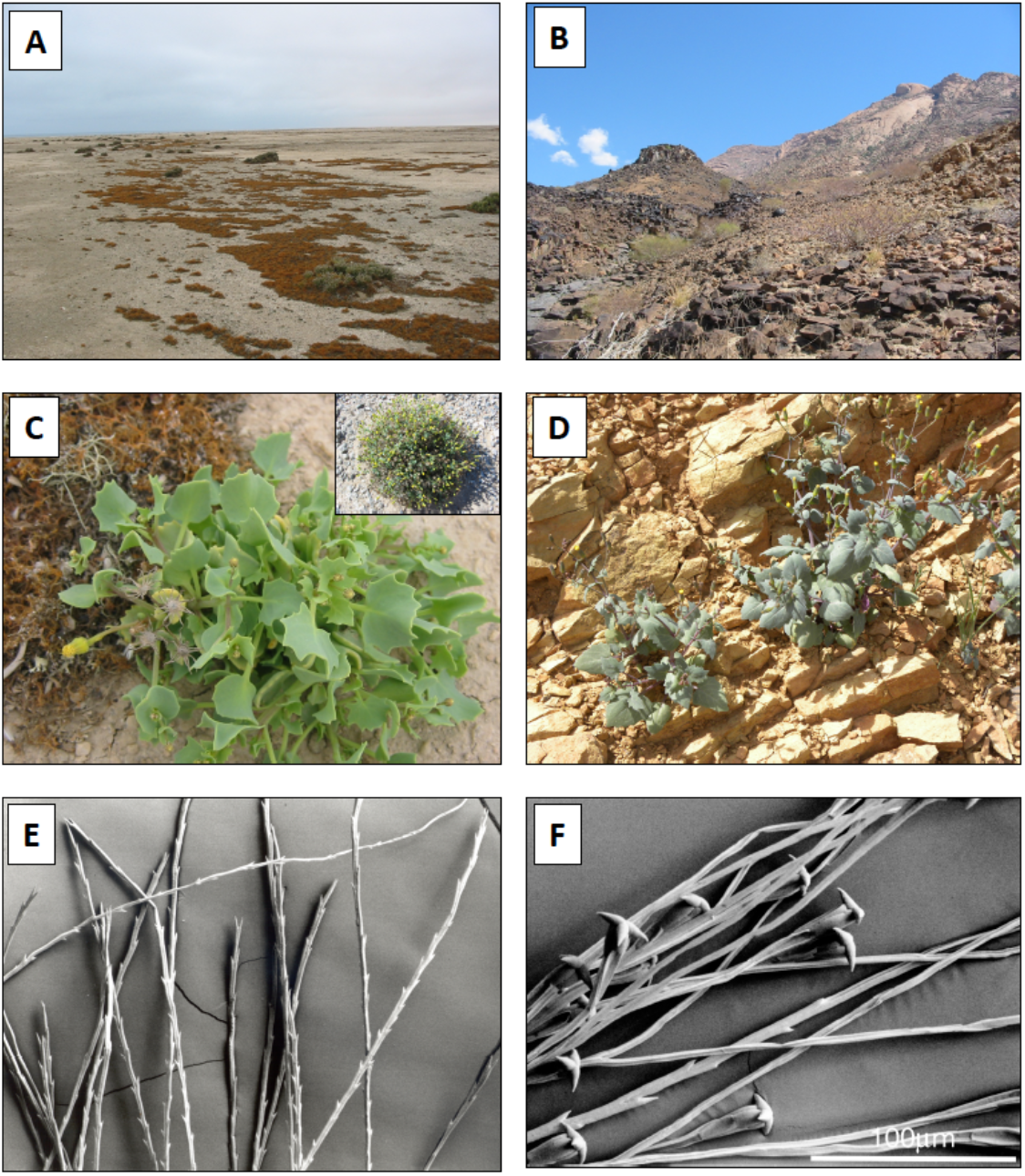
Habitat, phenotypes and pappus hairs of *Senecio englerianus* and *S. flavus*. (A) Gravelly coastal habitat of *S. englerianus* in the hyper-arid Central Namib Desert (near Swakopmund, pop. 3; see Table 1), associated with fruticose lichens (e.g. *Teloschistes capensis*) and Pencil Bush (*Arthraerua leubnitziae*, Amaranthaceae; see also Jürgens et al. 2013). (B) Rocky outcrop habitat of *S. flavus* in the (semi-)arid Nama-Karoo (near Keetmanshoop, c. 500 km south of Windhoek); here, the species is known to occur (B. Nordenstam, pers. comm.; see also Figure 1(B)), but was not found there during field work in April 2005. (C) Individual of *S. englerianus* near Swakopmund (pop. 3); the insert shows a large, cushion-forming individual from the same area (pop. 4) with more than a single year’s growth. (D) Individual of *S. flavus* from Southwest Morocco/Anti-Atlas. (E) Non-fluked (‘typical’) pappus hairs of *S. englerianus*, with forward-pointing teeth. (F) Connate-fluked pappus hairs of *S. flavus*, with grappling-hook-like appendages (reproduced from Coleman et al. 2003, with permission from Wiley & Sons, UK, 2 December 2021). Photographs by Joseph J. Milton (A–C), Jean-Paul Peltier (D; CC BY-NC 4.0), and Max Coleman (E, F).

**Table 1.** Locality information of populations and accessions of Senecio englerianus and S. flavus used for phylogenetic (phyl), morphometric (morph) and experimental-crossing analyses (marked with an asterisk). GenBank accession numbers are provided for all previously published and newly generated ITS sequences (the latter in italics). [XXX indicates that GenBank numbers will be added].

| Species | Population number/code<sup>a</sup> | Country, locality | Latitude | Longitude | Elev (m) | N (phyl) | N (morph) | Collector(s) (herbarium code) | GenBank accession<sup>b</sup> |
| --- | --- | --- | --- | --- | --- | --- | --- | --- | --- |
| *S. englerianus* | 1 | Namibia, Huab River | NA | NA | NA | 1 | – | Muller & Loutit 1164 (M) | AF457417<sup>s,p</sup> |
| *S. englerianus* | 2 * | Namibia, Swakopmund, road between Walvis Bay and Rooikop Airport, nr. quarry on the D road off the C14 road | 22° 40' S | 14° 33' E | 60 | 1 | 6 | JJM 110.1 (E) | XXX<sup>p</sup> |
| *S. englerianus* | 3 * | Namibia, Swakopmund, lichen-desert between Swakopmund and Henties Bay, c. 40 km from Swakopmund. | 22° 21' S | 14° 26' E | 24 | 1 | 7 | JJM 111.1 (E) | XXX<sup>p</sup> |
| *S. englerianus* | 4 | Namibia, Swakopmund, Henties Bay, in dried up mouth of Omaruru River | 22° 07' S | 14° 17' E | 3 | 1 | – | JJM 113.1 (E) | XXX<sup>p</sup> |
| *S. flavus* | MA | Namibia, Maltahöhe | 24° 29' S | 16° 58' E | 1,346 | 1 | – | Merxmüller & Giess 28206 (M) | AF457416<sup>s,p</sup> |
| *S. flavus* | TA (SF) * | Morocco, Souss-Massa Region, Tafraoute, roadside on outskirts of town | 29° 43' N | 08° 58' W | 1,000 | 2 | 5 | AA & RJA s.n. (STA) | TA1: XXX<sup>p</sup><br>TA2: XXX<sup>p</sup> |
| *S. flavus* | AS (SF751) * | Morocco, Marrakesh-Safi Region, 21 km from Asni (near Imgdal) | 31° 07' N | 08° 07' W | 1,568 | 1 | – | Ait Lafkih et al. 751 (RNG) | AF457414<sup>s</sup> |
| *S. flavus* | FU (SF2615) | Spain, Canary Isl., Fuerteventura, Puerto de la Peña/Ajuy | 28° 24' N | 14° 09' W | 97 | 1 | 1 | Nydegger 26145 (RNG) | AF457415<sup>p</sup> |
| *S. flavus* | SI | Egypt, Sinai, road from Sharm-El Sheik to Dahab | 28° 30' N (approx.) | 34° 30' E (approx.) | 0 | 1 | – | Coleman 10/00 (E) | AJ400782<sup>p</sup> |

Abbreviations: N, number of individuals used for phylogenetic (phyl) and morphometric (morph) analyses, respectively; NA = not available; Elev, elevation (in meters above sea level); collector abbreviations: AA, Anna Abbott; JJM, Joseph J. Milton; RJA, Richard J. Abbott; herbarium abbreviations: E, Royal Botanic Garden Edinburgh Herbarium; M, Munich Herbarium (Botanische Staatssammlung München); RNG, University of Reading Herbarium; STA, University of St Andrews Herbarium.
<sup>a</sup> Population codes of *S. flavus* given in parentheses refer to those in Milton (2009).
<sup>b</sup> Sequences included in the BEAST analysis of Senecioneae (Figures 3 and S1) and ML analysis of the ‘petiolate’ clade of *S. englerianus*/*S. flavus* (Figure S2) are indicated by superscripts ‘S’ and ‘P’, respectively.

These diploid herbs (each 2*n* = 2x = 20; Coleman et al. 2001) are morphologically similar, and share characteristic petiolate leaves. However, *S. englerianus* is a short-lived perennial (Merxmüller 1967; Jürgens et al. 2021; JJ Milton, pers. obs.), has smaller, more succulent leaves and larger capitula, and is dimorphic for capitulum-type (mostly non-radiate, occasionally radiate; Jürgens et al. 2021), whereas *S. flavus* is an annual herb with non-radiate capitula throughout its range (Alexander 1979; Coleman et al. 2001; e-Flora of South Africa 2018; Figures 2(C) and (D)). Furthermore, in *S. englerianus*, the pappus hairs have forward-pointing teeth, as typical for the genus (Drury and Watson 1996), and detach easily from the achene (Figure 2(E)); by contrast, *S. flavus* has additional, so-called ‘connate-fluked’ pappus hairs with grappling-hook-like appendages (accounting for about 1/3 of the entire pappus; Figure 2(F)), and which might enhance the species’ dispersal capacity (Coleman et al. 2003). Finally, *S. flavus* is fully self-compatible (Liston et al. 1989; Coleman et al. 2003), with an estimated selfed seed set (mean ± S.E.) of 74.2 ± 4.2% (N = 40), while *S. englerianus* fails to set seed when left to self (JJ Milton, unpublished data).

Based on isozyme data and biogeographical considerations, Liston et al. (1989) hypothesized that *S. flavus* originated in Southwest Africa and expanded its range to North Africa, possibly via an African Dry-Corridor (ADC) during periods of Pleistocene aridification (e.g. van Zinderen Bakker 1975; reviewed in Neumann and Scott 2018). Subsequent phylogenies of Senecioneae, inferred from internal transcribed spacer (ITS) regions of nuclear ribosomal (nr) DNA, indeed suggested that the closest relative of *S. flavus* is *S. englerianus* (Coleman et al. 2003; Pelser et al. 2007). In both studies, this highly supported and long-branched ‘petiolate’ clade (*sensu* Coleman et al. 2003) appeared only distantly related to sect. *Senecio* or South African species. Furthermore, Coleman et al. (2003) found no nucleotide differences between *S. englerianus* and Namibian *S. flavus,* and (roughly) dated the split between the latter and North African *S. flavus* (one nucleotide difference) to the Late Pleistocene, c. 0.15 million years ago (Mya).

Overall, *S. englerianus* and *S. favus* provide an excellent system for evaluating the hypothesis of budding speciation in a putative pair of sister species that occur in parapatry in an isolated arid ecosystem, including the identification and characterization of RI barriers. In addition, by using palaeodistribution modelling (Phillips et al. 2017), the disjunct distribution of *S. flavus* allows for testing the ADC hypothesis of an ‘arid track’ promoting plant migration from Southwest to North Africa during the Pleistocene. Accordingly, the specific aims of the present study were as follows. First, to re-examine the putative sister relationship of *S. englerianu*s and *S. flavus* (Coleman et al. 2003; Pelser et al. 2007) and, in particular, to estimate their divergence time, based on a more extensive taxon sampling (especially of Southwest and South African species), the inclusion of multiple intraspecific accessions of *S. englerianus*/*S. flavus*, and modern techniques in molecular dating. Second, to determine the extent of environmental niche overlap and differentiation between the two species in Namibia, and their morphological divergence. Third, to evaluate differences in mating system based on pollen counts (pollen/ovule ratios; Cruden 1977) and the genetic basis of ‘intrinsic’ postzygotic isolation (based on F_2_ pollen fertility) and differences in pappus type between *S. englerianus* and *S. flavus*. Fourth, to test for a potential role of the ADC in promoting the hypothesized northward expansion of *S. flavus*, using palaeodistribution modelling.

## Materials and methods

### Field work in Namibia and experimental plant material

In April 2005, field work was conducted by one of us (J.J.M.) for two weeks in western and southern Namibia. This seasonal timing (at the end of the summer-rain period) was considered suitable based on phenological dates derived from herbarium specimens (personal collection Bertil Nordenstam/Swedish Museum of Natural History). Seed and silica-dried leaf material was collected from three coastal populations of *S. englerianus* in the Central Namib Desert (Table 1), but despite intensive searching, no populations of *S. flavus* were found. Notably, in Namibia, this (semi-)desert annual is reported to flower in response to rainfall, being “very common in years of good rain and apparently very rare in drier years”; c.f. Foden et al. 2011). Instead, for comparative cultivation and crossing experiments, individuals of *S. flavus* were grown from seed material collected previously from Canary Island and Moroccan populations (see Table 1 and further below).

### *Assembly of geo-referenced locality records for* S. englerianus *and* S. flavus

To illustrate the species’ distributions at the contintental-wide scale and in Southwest Africa (Figure 1), we retrieved geo-referenced locality records for *S. englerianus* vs. *S. flavus* from various sources, including (1) the Global Biodiversity Information Facility [GBIF.org; *S. englerianus*: GBIF.org (5 November 2020) GBIF Occurrence Download https://doi.org/10.15468/dl.4be9tr; *S. flavus*: GBIF.org (9 November 2020) GBIF Occurrence Download https://doi.org/10.15468/dl.gj8yzk]; (2) the Plants of Namibia Datbase/BRAHMS online (Craven and Kolberg 2021; https://herbaria.plants.ox.ac.uk/bol/namibia); (3) iNaturalist (inaturalist.org); (4) published sources (Coleman et al. 2003); (5) the personal collection of B. Nordenstam (see above); and (6) own field work in Namibia (only *S. englerianus*; full data available on request). Duplicate occurrence records per species and GBIF-records outside known taxon ranges were filtered out. Maps were generated in QGIS v3.8 (QGIS Development Team 2019) after retaining 30 locality records of *S. englerianus* and 150 of *S. flavus* [Southwest Africa (i.e. Namibia/northwestern South Africa): 33; eastern South Africa (KwaZulu-Natal): 1; Canary Islands (Fuerteventura), North Africa (Morocco to Middle East) and Southeast Spain: 116]. We also used these records for niche analyses and species distribution modelling (see below).

### Phylogenetic analyses of Senecioneae and the ‘petiolate’ clade

To estimate the stem and crown group ages of the ‘petiolate’ clade of *S. englerianus*/*S. flavus*, and its phylogenetic position within Senecioneae (c. 150 genera, > 3,000 spp.; Pelser et al. 2007), we first retrieved 371 ITS sequences (c. 340 spp.) from GenBank (e.g. Panero et al. 1999; Pelser et al. 2007, 2010; Kandziora et al. 2017), representing (1) one acession of *S. englerianus* (pop. 1) and three of *S. flavus* [Namibia, Morocco (AS), Fuerteventura]; (2) all major clades of *Senecio* s.str. (e.g. ‘*vulgaris*’, ‘*windhoekensis*’, ‘*suaveolens*’; *sensu* Kandziora et al. 2017); (3) 27 genera of subtribe Senecioninae (e.g. *Arrhenechthites*, *Crassocephalum*, *Curio*, *Erechtites*, *Jacobaea*, *Kleinia*); (4) two genera of subtribe Othonnineae (*Euryops*, *Othonna*); and (5) one species of tribe Athroismeae (*Anisopappus pinnatifidus*). We further complemented this ‘Senecioneae’ ITS dataset with 33 newly generated sequences of sub-Saharan (mostly Namibian and/or South African) species of *Senecio* (31 spp./32 accessions) and *Kleinia crassulaefolia* (South Africa). Respective leaf samples were obtained from field work by one of us (J.J.M.) in South Africa (September 2004 and 2005; 28 spp.), the living collection of the Royal Botanic Garden Edinburgh (RBGE; 3 spp.), and a personal herbarium (Michael Möller, RBGE). Details of DNA extraction, ITS amplification, sequencing and cloning (4 spp.) can be found in the Supplementary Information (see Appendix 1). A full list of taxa included in the phylogenetic analysis of Senecioneae (403 accessions, c. 363 spp.), together with accession numbers, information on species distributions and locality sources of newly sequenced ITS samples, is provided in Appendix 2 (Tables A2.1–3).

**Table 2.**
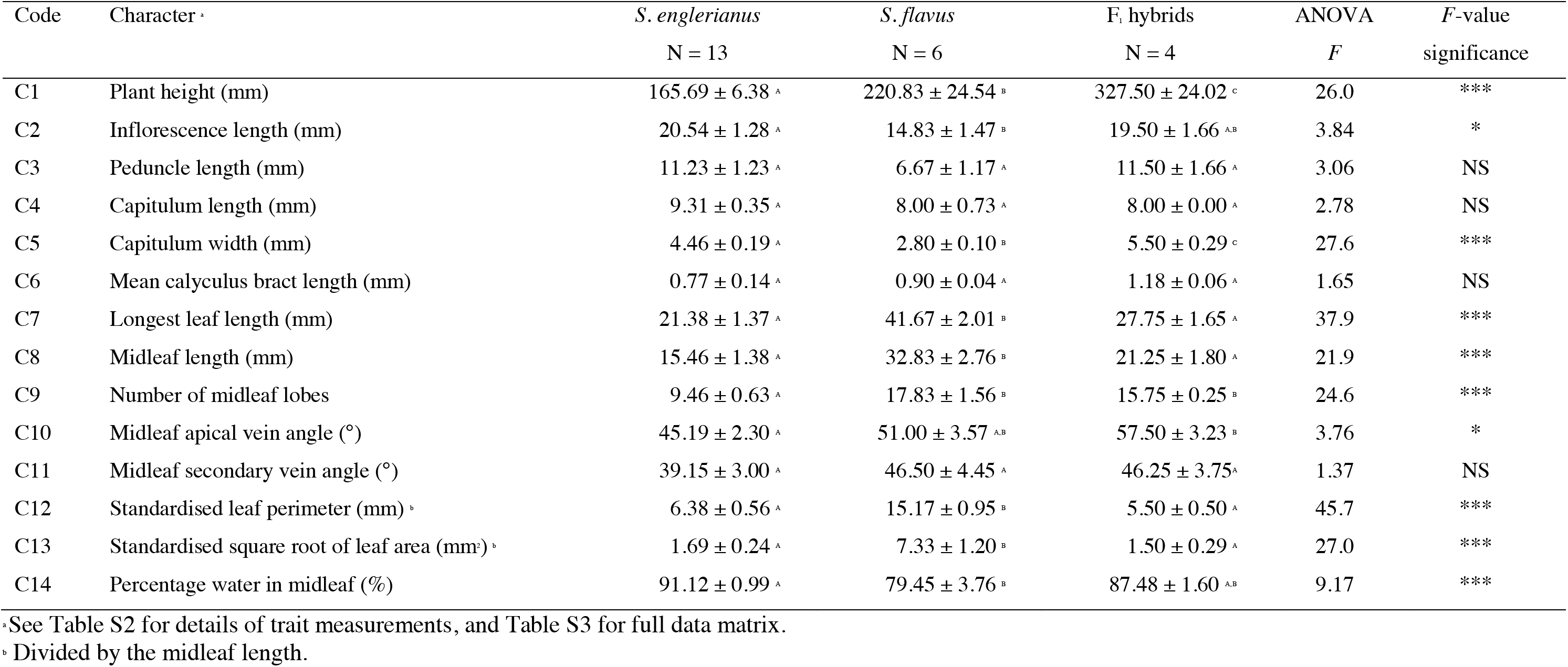
Means (± S.E.) of 14 trait variables (C1–C14) measured on 13 individuals of Senecio englerianus (six from pop. 2, seven from pop. 3), six individuals of S. flavus [five from Morocco (TA), one from Fuerteventura (FU)] and four F_1_ hybrids created by crossing the two species (i.e. two individuals each of crosses AS.1 x 3.29 and TA.5 x 2.2; see text). Shared superscript letters indicate no significant difference at the 5% level using Tukey’s multiple comparisons HSD (honestly significant difference) test. One-way analysis of variance (ANOVA) F-value significance: * P < 0.05; ** P ≤ 0.01; *** P ≤ 0.001; NS, not significant. Note, non-normally distributed measurements of size (C1–C8) were natural log-transformed, and C14 was arcsine transformed.

Sequences of this Senecioneae dataset were assembled and aligned in GENEIOUS v8.1.9 (http://www.geneious.com; data matrix available on request). Bayesian searches for tree topologies and node ages were conducted in BEAST v1.7.5 (Drummond et al. 2012) with the GTR+G substitution model and an uncorrelated lognormal relaxed clock (Drummond et al. 2006). The tree speciation prior followed a Yule process. For calibration, we assumed a normal distribution of the ITS substitution rate of Asteraceae with a mean (± S.D.) of 4.59 ± 2.42 x 10^-9^ substitutions/site/year (Kay et al. 2006). Three independent Markov chain Monte Carlo (MCMC) simulations were run for 10^8^ generations, sampling every 10^4^ generations. The three runs were combined after removing 67% of the trees; the remainder was used to construct a maximum clade credibility (MCC) chronogram with clade posterior probabilities (PP), mean node ages and 95% highest posterior density (HPD) intervals. The full MCC topology was also vizualized as ‘circle tree’ using PHYTOOLS v0.7–90 (Revell 2012).

To further clarify relationships within the ‘petiolate’ clade, we assembled all GenBank ITS sequences of *S. englerianus* (N = 1) and *S. flavus* (N = 4) with six newly generated ones [one each per pops. 2–4 vs. two from Morocco (TA); see Table 1], plus all GenBank accessions of the *Arrhenechthites*–*Dendrocacalia* clade (4 spp.), as putative sister group based on the Senecioneae phylogeny (see Results). We then used RAxML (Stamatakis 2014) from the raxmlGUI v2.0 platform (Edler et al. 2021) to construct a maximum likelihood (ML) tree under the default GTR+G model. The best-fit tree was rooted with *Arrhenechthites novoguineensis* and annotated with bootstrap support (BS) values based on 1000 replicates.

### Niche overlap and divergence analyses

We used the environmental-(E)-space-based framework of the R package HUMBOLDT (Brown and Carnaval 2019) to study niche overlap and divergence between *S. englerianus* and *S. flavus* in Southwest Africa, based on our 63 presence records for this region (N = 30 vs. 33; Figure 1(B); i.e. excluding one *S. flavus* outlier record from eastern South Africa; Figure 1(A)). We first retrieved 19 bioclimatic variables (bio1–19) at 2.5 arc-minutes resolution for current conditions (1950–2000) from WORLDCLIM v1.4 (Hijmans et al. 2005; http://www.worldclim.org/) and downloaded elevation data from WORLDCLIM v2.1 (Fick and Hijmans 2017; https://www.worldclim.org/data/worldclim21.html). We subsequently removed highly correlated climatic variables (Pearson’s *r* > |0.9|) in R using ENMTOOLS v1.4.4 (Warren et al. 2010) to minimize model overfitting. Based on altitude and 12 climatic variables retained (i.e. bio1, 3, 6, 7, 9–11, 15–19; see Table S1 for code identification), we conducted a principal component analysis (PCA), along with kernel density functions, to visualize the species’ environmental niche overlap in two dimensions.

In addition, we performed two niche similarity tests implemented in HUMBOLDT, i.e. ‘niche overlap test’ (NOT) and ‘niche divergence test’ (NDT), each of them including the calculation of equivalency (*E*) and background (*B*) statistics using 500 replicates. In brief, NOT determines how equivalent (or dissimilar) the occupied niches of the two species are, given their total E-space, while NDT determines significant differences in the species’ *shared* E-space (Brown and Carnaval 2019; see also Habibzadeh et al. 2021). In either instance, the *E* statistic compares the observed value of Schoener’s (1968) *D* metric between the two species, ranging from 0 (no niche overlap) to 1 (complete overlap), with a null distribution obtained from the species’ resampled and spatially shifted locality records. In contrast, the *B* statistic compares the interspecific *D* value observed to differences between *S. englerianus* and the resampled and reshuffled occurrences of Southwest African *S. flavus*, and vice versa (denoted *B*_flav–>engl_ and *B*_engl–>flav_, respectively). The *B* statistic evaluates if two species are more different than would be expected given the underlying environmental differences between the regions in which they occur. However, regardless of the *B* statistic, a significant *E* statistic in both NOT and NDT provides strong evidence that the species’ niches are not only different but have also diverged. The same applies if very low *D* values are observed in both tests and NOT is significant (see Brown and Carnaval 2019, for tabulation of interpreting other results).

### Plant cultivation and controlled crossing experiments

Plants were raised from fruits (achenes) collected from different mother plants in each of three populations of *S. englerianus* (pops. 2–4) and *S. flavus* [Morocco (AS, TA), Fuerteventura (FU); Table 1]. After germinating achenes on damp filter paper, seedlings were transplanted to pots (7.6 cm diameter) containing a 3 : 1 mixture of Levington’s M2 compost to gravel. Plants were arranged in a randomized block design and raised at ambient temperature in a greenhouse with natural light, supplemented by 400 W mercury vapour lamps with photoperiod set to 16 h. In addition, experimental crosses were made between one individual each of *S. englerianus* populations 2 and 3 (specimens 2.2 and 3.29) and one individual each of Moroccan *S. flavus* from AS and TA (AS.1 and TA.5). In all crosses, individuals of *S. flavus* were used as maternal parent and subjected to Alexander’s (1979) emasculation technique, resulting in two self-fertile F_1_ generations (AS.1 x 3.29 and TA.5 x 2.2). One F_1_ individual of the AS.1 x 3.29 cross was further used for the production of an F_2_ generation through self-pollination (bagging). All F_1_ and F_2_ plants were raised in the greenhouse as described above.

### Quantitative trait analyses

At full anthesis of the apical capitulum, 14 quantitative characters (C1–C14) were scored for 13 individuals *S. englerianus* (six from pop. 2 and seven from pop. 3), six individuals of *S. flavus* (five from TA and one from FU), and four F_1_ hybrids created by crossing the two species (i.e. two individuals each of crosses AS.1 x 3.29 and TA.5 x 2.2; see above). Following Lowe and Abbott (2000), the morphological traits measured (C1–C13) included: plant height, inflorescence and peduncle length, capitulum length and width, and various descriptors of leaf size and shape (for details, see Table S2). In addition, we measured leaf water content (C14), a physiological proxy for tissue succulence (Fradera-Soler et al. 2021), by weighing midleaves, drying them overnight in an oven and then reweighing.

For statistical analysis, non-normally distributed measures of size (C1–C8) were natural log-transformed; leaf perimeter (C12) and square root of leaf area (C13) were divided by midleaf length to standardize; and percentages of leaf water content (C14) were arcsine transformed. The full 23-individuals x 14-character matrix (Table S3) was subjected to one-way analysis of variance (ANOVA) and Tukey’s HSD (honestly significant difference) post-hoc test using the R package DPLYR v1.0.6 (Wickham et al. 2021) to test for significant inequality of character means among groups (both species and the F_1_ plants) and to detect which means differed significantly from each other. Finally, the data matrix was subjected to a PCA in R v3.6.2 (R Development Core Team 2017) and visualized in two dimensions using GGBIPLOT v0.55 (Vu 2011). To complement traditional PCA loadings, we also calculated the contribution of each variable (in %) to each ordinated axis (PC1, PC2), using the R package FACTOEXTRA v1.0.5 (Kassambara and Mundt 2017).

### Estimates of pollen number and pollen fertility in parental, F_1_ and F_2_ plants

Pollen number and fertility were estimated for one individual each of *S. englerianus* (3.29), Moroccan *S. flavus* (AS.1), and the F_1_ (AS.1 x 3.29), as well as 32 individuals of the F_2_ generation, produced by self-pollination of this F_1_ hybrid. Total pollen counts were made by means of light microscopy, using 2–5 florets per capitulum and one or two capitula per individual. Concomitantly, pollen fertilities were estimated by staining viable pollen with aceto- carmine. Mean pollen numbers and mean percentages of pollen fertility were calculated for parental species, the F_1_ hybrid and the F_2_ generation (for the full data matrix, see Table S4). Resulting frequency distributions of F_2_ pollen counts and fertility were compiled in histograms and examined for normality using Shapiro-Wilks (*W*) test in R, with *P* > 0.05 indicating a normal distribution. As the F_2_ histogram for pollen fertility indicated a bimodal pattern (see Results), we applied the Expectation Maximization (EM) algorithm (Dempster et al. 1977) to fit two Gaussian (normal) distributions to this dataset, using the R package MIXTOOLS v1.2.0 (Young et al. 2020). Frequency estimates of each class and distribution were subjected to Mendelian ratio/Chi-square analyses to evaluate the genetic basis of reduced F_2_ hybrid fertility (viz. hybrid incompatibility; e.g. Johnson 2008). Note, with two peaks in an F_2_ phenotype distribution, four possible Mendelian ratios are expected (Griffiths et al. 2000): 3 : 1 (one full dominant gene); 9 : 7 (two duplicate recessive epistatic genes); 13 : 3 (two recessive-dominant epistatic genes); and 15 : 1 (two duplicate dominant epistatic genes).

### Genetic analysis of pappus type

To examine the genetic basis of present vs. absent connate-fluked (CF) pappus hairs in *S. englerianus* vs. *S. flavus* (Coleman et al. 2003; Figures 2(E) vs. (F)), we examined the segregation of this trait in the same F_2_ family as described above for the analysis of pollen number and fertility. Accordingly, one individual each of the two species, their self-fertile F_1_ hybrid (AS.1 x 3.29), and 87 individuals of the F_2_ generation were examined for pappus type by light microscopy. Observed morph frequencies were evaluated by Chi-square analysis to evaluate their adequacy with a single gene model, as often reported for presence/absence dimorphisms (e.g. Gottlieb 1984; Lankinen 2009).

### *Present and past distribution modelling of* S. englerianus *and* S. flavus

Based on our locality records of *S. englerianus* (N = 30) and Southwest African *S. flavus* (N = 33), ecological niche models (ENMs) were generated for each dataset in MAXENT v3.4.1 (Phillips et al. 2017) for the present, and two past periods: the humid mid-Holocene Climate Optimum (HCO; c. 6 kyr ago) and the arid Last Glacial Maximum (LGM; ca 22 kyr ago). This allowed us to determine whether the ranges of the two species in Southwest Africa were also continuous (para- or sympatric) in the past (rather than allopatric), and whether *S. flavus* could have reached North Africa via the ADC during periods of aridification as hypothsized by Liston et al. (1989). For each dataset, the current distribution (1950–2000) was modelled using 11 bioclimatic layers at 2.5 arc-minutes resolution (bio2, 7–11, 15–19; Table S1) after removing highly correlated variables (*r* > |0.9|). Each model was then projected on to palaeoclimatic datasets simulated by the Model for Interdisciplinary Research on Climate (MIROC) for, respectively, the HCO (MIROC-ESM; http://www.worldclim.org/) and the LGM (MIROC- ESM; http://www.worldclim.org/). Present-day distributions were modelled 10 times, using different subsets of 75% of the localities to train the model. Model accuracy was evaluated for each run, and for both training and testing data, using the Area Under the Curve (AUC) of the Receiver Operating Characteristic (ROC) plot. For comparison, we also generated present and past ENMs for all 150 records of *S. flavus* from across the species’ range.

## Results

### Phylogeny reconstruction, molecular dating and intra-clade relationships

Our BEAST-derived MCC chronogram of Senecioneae (Figure 3) recovered representative accessions of *S. englerianus* (pop.1) and *S. flavus* [Namibia, Morocco (AS), Fuerteventura] as a highly supported clade (PP = 1.0; see Figure S1 for the full tree topology). However, this apparently highly isolated ‘petiolate clade (*sensu* Coleman et al. 2003) essentially formed a polytomy with three other clades (each PP = 1.0), namely *Senecio* s.str., C*rassocephalum*– *Erechtitis* and *Arrhenechthites*–*Dendrocacalia* (including *S. thapsoides*). We estimated the stem age of the *S. englerianus*/*S. flavus* clade to the early Late Miocene, c. 10.91 Mya, albeit with large 95% HPDs (23.14–3.26 Mya), and its crown age to the Early Pleistocene, c. 1.47 (2.72–0.11) Mya (see Table S5 for divergence dates of other major clades and lineages).

**Figure 3.**
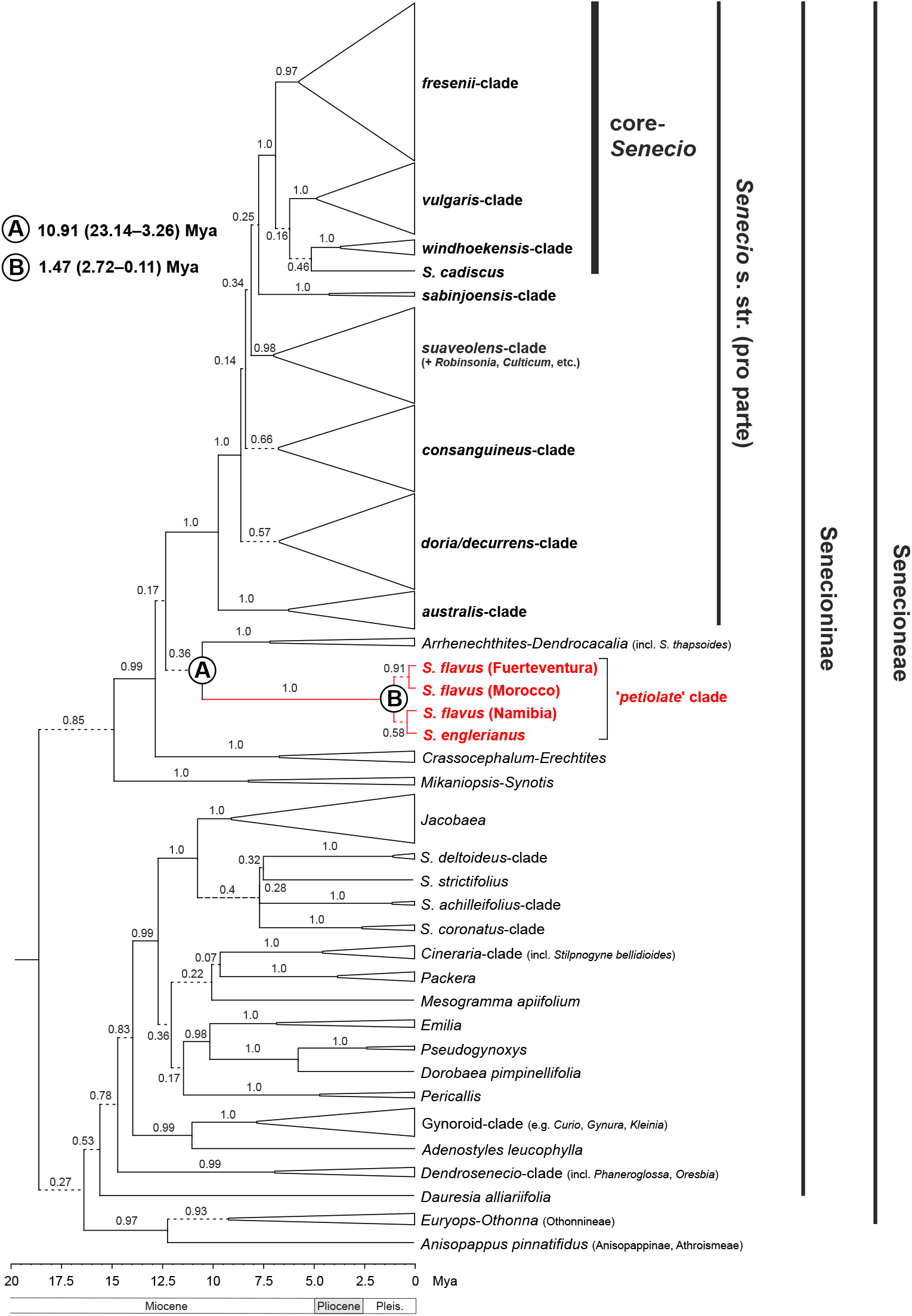
(A) BEAST-derived maximum clade credibility (MCC) chronogram of Senecioneae, plus one member of Athroismeae (*Anisopappus pinnatifidus*), based on ITS sequences, with collapsed major clades. Numbers above branches indicate Bayesian posterior probability (PP) values. Branches in dashed lines indicate PP ≤ 0.95. Designation of clades within *Senecio* s.str. largely follows Kandziora et al. (2017). Mean estimates of stem (A) and crown (B) ages for the ‘petiolate’ clade (in million years ago, Mya) are indicated, together with 95% highest posterior density (HPD) intervals (in parentheses). Respective time estimates and 95% HPDs for all other major clades and single species lineages are summarized in Table S5. Pleis, Pleistocene. See Figure S1 for the full MCC chronogram of all 403 accessions (c. 363 spp.).

However, relationships within this ‘petiolate’ clade largely remained unresolved (Figure 3), and even after subjecting all available ITS sequences of each species (10 in total) to ML analysis, with species of the *Arrhenechthites*–*Dendrocacalia* clade serving as outgroup (see Figure S2). Nonetheless, while all accessions of *Senecio englerianus* (N = 4) and the single one of Namibian *S. flavus* essentially collapsed into a polytomy, all northern material of *S. flavus* [Morocco (AS, TA), Fuertventura, Egypt; N = 5] formed a weakly supported cluster (BS = 57%) due to a single, shared derived nucleotide substitution (T to C transition).

### *Niche differentiation between* S. englerianus *and* S. flavus *in Southwest Africa*

The HUMBOLDT-derived PCA biplot and associated density plots (Figure 4(A)) revealed limited environmental niche overlap (purple) between *S. englerianus* (blue, higher scores) and *S. flavus* (red, lower scores) in Southwest Africa, largely due to differences along the first component (PC1), explaining 52.8% of the total variance (PC2: 21.3%). Considering the corresponding correlation circle of 13 environmental variables (Figure 4(B)), PC1 was more or less positively correlated with isothermality (bio3), minimum temperature of coldest month (bio6), measures of quarterly temperature (bio9, 11), and precipitation seasonality (bio15), and negatively loaded for temperature annual range (bio7), indices of quarterly precipitation (bio16–19), and altitude.

**Figure 4.**
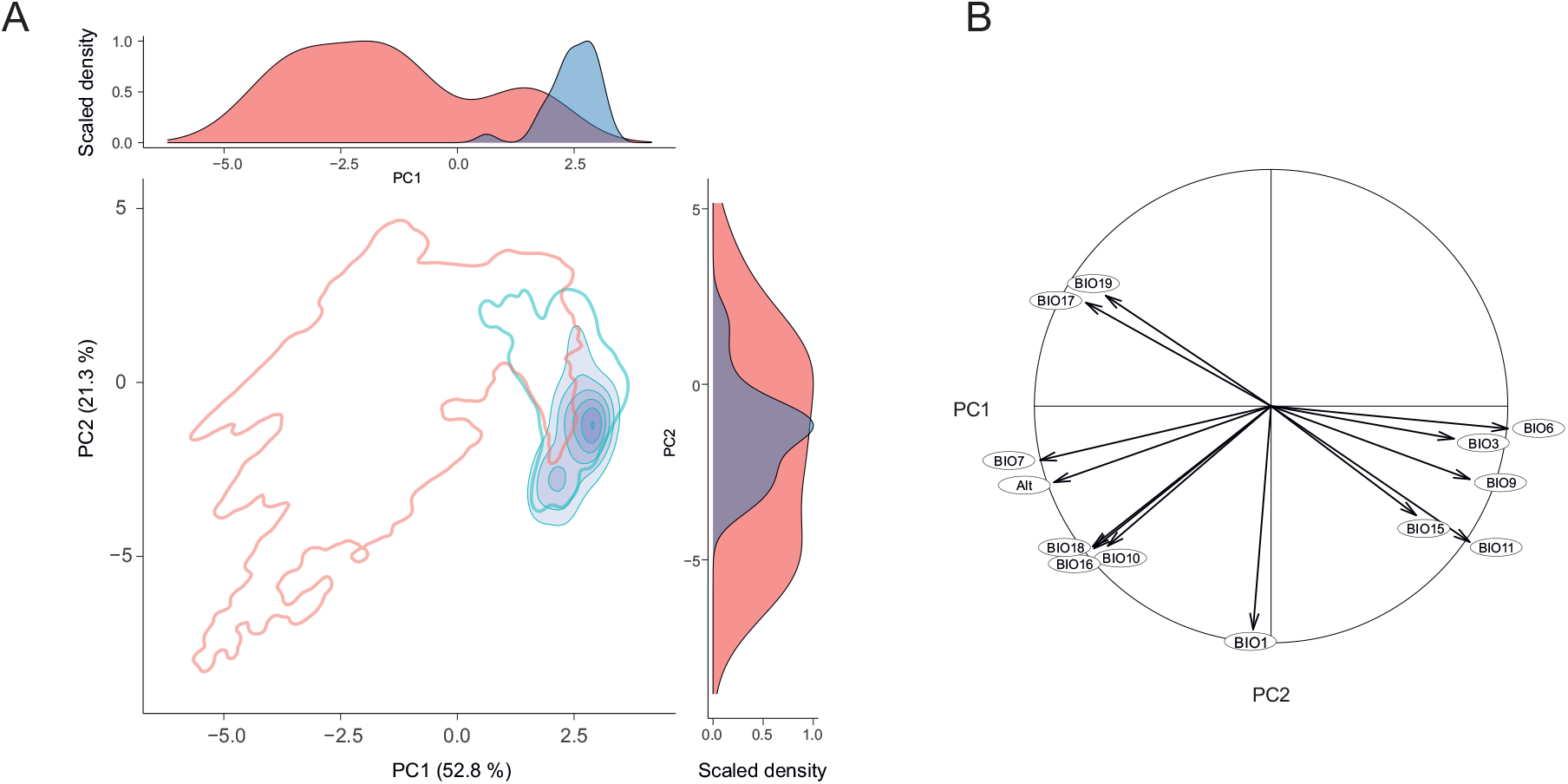
(A) HUMBOLDT-derived principal component analysis (PCA) biplot, along with density plots for each component (PC1, PC2), depicting the extent of environmental niche overlap (purple) between *Senecio englerianus* (blue) and *S. flavus* (red) in Southwest Africa. Percentages of total variance explained by each component are shown in parentheses. Filled kernel isopleths present kernel densities from 1–100%. Empty kernel isopleths represent the 1% density isopleths of the environments (not species) outside boundaries of environmental (E) space. (B) Corresponding correlation circle of 12 bioclimatic variables (see Table S1 for code identification) and altitude (Alt). The length and directionality of arrows (vectors) illustrate, respectively, the strength of contribution (loading) and sign of correlation (positive, negative) of each variable on the two components (PC1: x-axis; PC2: y-axis). Variables closer to each other are thus more correlated on a given component. Analyses were conducted based on 63 locality records (*S. englerianus*: N = 30; *S. flavus*: N = 33; see also Figure 1(B)).

Hence, relative to *S. englerianus*, *S. flavus* tends to occur at higher elevation sites, mostly in the Nama-Karoo (Figure 1(B)), which experience stronger fluctuations in annual temperature (bio3, 7) as well as higher and more predictable seasonal or annual rainfall (bio15– 19). For the niche overlap test (NOT), Schoener’s *D* value in the *E* statistic was highly significant and near-zero (*D* = 0.003, *P* = 0.002), while the two *B* statistics were non-significant (*B*_flav–>engl_: *P* = 0.375; *B*_engl–>flav_: *P* = 0.582). The *E* statistic of the niche divergence test (NDT) yielded a *D* value of zero, albeit without significance (*P* = 0.21), while the two *B* statistics were significant (*B*_flav–>engl_: *P* = 0.003; *B*_engl–>flav_: *P* = 0.012). The non-significant *D* value in the NDT could be due to lack of statictical power viz. insufficient records in the species’ limited area of overlap (i.e. Central Namib Desert and hinterland; Figure 1(B)), or might indicate real absence of shared available E-space. Regardless, a significant NOT combined with (near-) zero *D* values in both NOT and NDT provides strong evidence that the environmental spaces occupied by the two species in Southwest Africa are not only significantly different (i.e. non-equivalent) but truly reflect divergent niche evolution (Brown and Carnaval 2019).

### Quantitative trait analyses

One-way ANOVA revealed significant (*P* ≤ 0.05) differences in means between *S. englerianus* (N = 13), *S. flavus* (N = 6; Morocco, Fuerteventura) and their F_1_ hybrids (N = 4) for 10 of the 14 characters recorded (Table 2). Tukey’s HSD tests showed significant differences between the means of *S. englerianus* and *S. flavus* for all of these characters except C10 (midleaf apical vein angle). Thus, relative to *S. flavus*, *S. englerianus* was shorter in height (C1), had longer inflorescences (C2) and wider capitula (C5), and produced smaller, more succulent leaves with fewer midleaf lobes (C7–C9, C12–14). For half of these characters (i.e. C2, C7–C9, C14), the F_1_ individuals showed an intermediate phenotype between both parental species. For two characters (height, capitulum width), the means of F_1_ plants were significantly greater than those of either parent species (Table 2).

In the corresponding PCA biplot (Figure 5), the first component (PC1), explaining 41.5% of the total variance, clearly separated *S. englerianus* and *S. flavus* into low- and high-score clusters, respectively, which largely reflected their differences in leaf size and shape (C7– C9, C12, C13) and leaf succulence (C14; see Table S2 for loadings and variable contributions). The F_1_ plants occupied an intermediate position between the parental clusters along PC1, yet closer to *S. englerianus*; however, they were largely separated from either parent along PC2 (17.9%), mainly because of their greater stature and higher values in other variables that were strongly negatively correlated with this second component [e.g. peduncle length (C3), calyculus bract length (C6), midleaf vein angles (C10/11); see Table S2].

**Figure 5.**
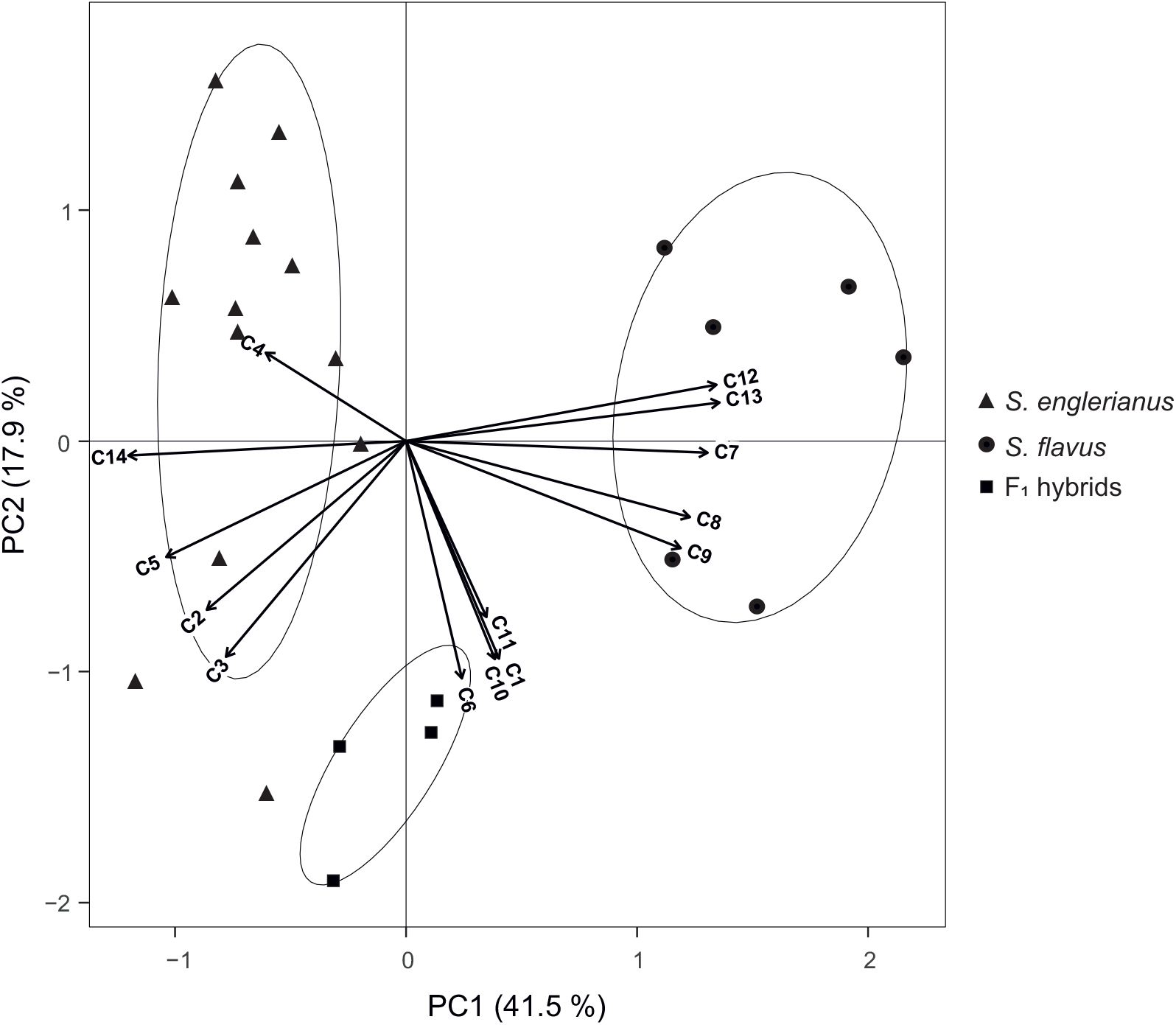
Principal component analysis (PCA) biplot of 14 quantitative trait variables (C1–C14) measured for *Senecio englerianus* (N = 13), *S. flavus* (N = 6; Morocco, Fuerteventura), and F_1_ hybrids (N = 4). Percentages of total variance explained by each component (PC1, PC2) are noted in parentheses. Ellipses were fitted to the PCA plane (rather than to the full multivariate data set) so as to include 95% of the points in each of the three groups. See Table 2 for identification of trait codes (C1– C14), and Table S2 for details of trait measurements as well as factor loadings and variable contributions (in %) for each component.

### Pollen number and fertilities of parental, F_1_ and F_2_ individuals

As summarized in Table 3, the *S. englerianus* parent plant (3.29) produced, on average, more than six times the amount of pollen grains per floret (mean ± s.d.: 3394.40 ± 153.15) than the *S. flavus* parent (AS.1) from Morocco (495.20 ± 46.95). However, in both individuals, pollen fertility was very high, particularly in *S. englerianus* (99.0% vs. 89.68%). The mean pollen count of their self-fertile F_1_ individual (AS.1 x 3.29) was approximately the same as in the *S. flavus* parent (409 ± 31.52 grains/floret), while pollen fertility was significantly reduced (59.3%). The mean pollen count of the F_2_ generation (N = 32) was intermediate to that of the parents (1825.35 ± 48.14 grains/floret), and mean pollen fertility (85.24%) was greater than that recorded for the F_1_ hybrid (Table 3).

**Table 3.**
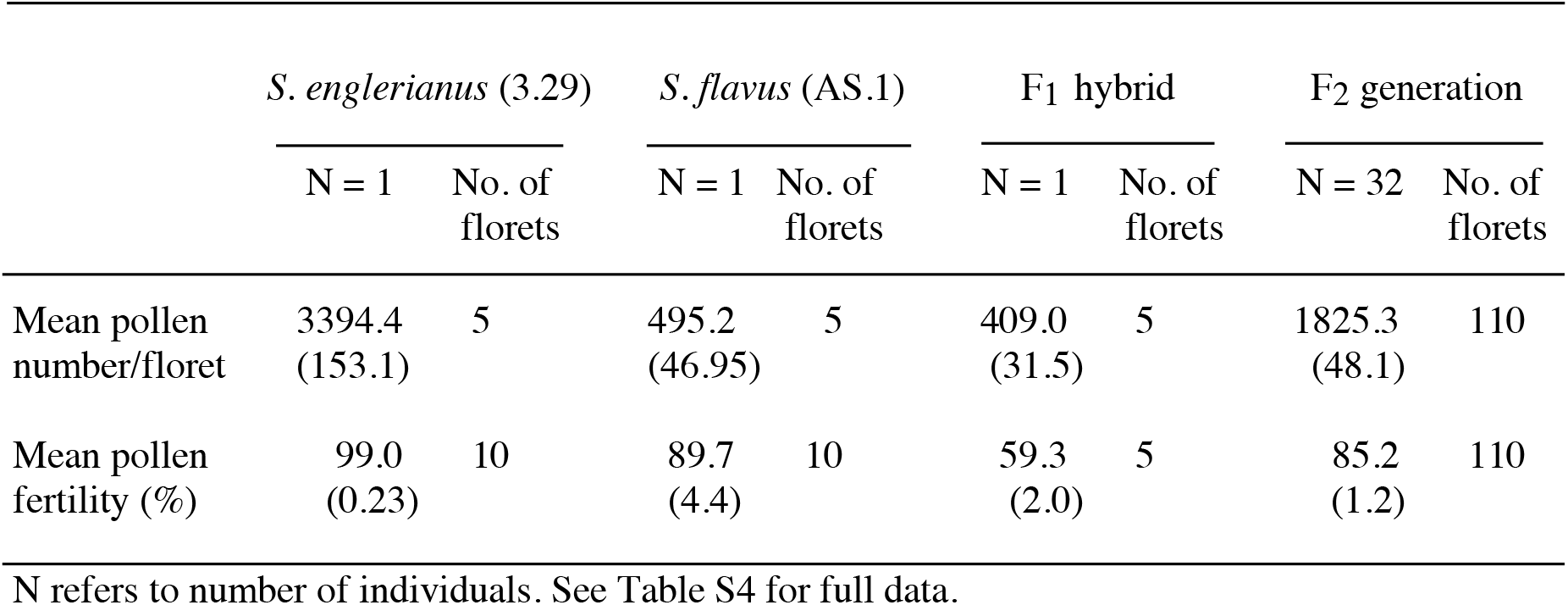
Mean estimates of pollen number (grains/floret) and percentage pollen fertility in *Senecio englerianus* (specimen 3.29), Moroccan *S. flavus* (AS.1), one of their F_1_ hybrids [AS.1 x 3.29 (1)], and 32 individuals of the F_2_ generation, produced after self-pollination of this F_1_ hybrid. Standard errors are shown in parentheses.

|  | *S. englerianus* (3.29) |  | *S. flavus* (AS.1) |  | F<sub>1</sub> hybrid |  | F<sub>2</sub> generation |  |
| --- | --- | --- | --- | --- | --- | --- | --- | --- |
|  | N = 1 | No. of florets | N = 1 | No. of florets | N = 1 | No. of florets | N = 32 | No. of florets |
| Mean pollen number/floret | 3394.4<br>(153.1) | 5 | 495.2<br>(46.95) | 5 | 409.0<br>(31.5) | 5 | 1825.3<br>(48.1) | 110 |
| Mean pollen fertility (%) | 99.0<br>(0.23) | 10 | 89.7<br>(4.4) | 10 | 59.3<br>(2.0) | 5 | 85.2<br>(1.2) | 110 |

N refers to number of individuals. See Table S4 for full data.

The pollen count data for the F_2_ generation (Figure 6(A)) fitted a normal distribution (Shapiro-Wilks’ *W* = 0.980, *P* = 0.781), likely reflecting recombination and segregation of alleles at several or many loci. Of the 32 F_2_ plants examined, 11 had mean pollen counts between 1750 and 2000 grains/floret, hence intermediate with respect to the values of the two parents (see above). Notably, none of the F_2_ individuals had mean pollen counts as low as either the *S. flavus* parent or the F_1_ hybrid, with the lowest values falling in the 1000–1250 grains/floret class. By contrast, the distribution of pollen fertilities for the F_2_ (Figure 6(B)) departed from normality (*W* = 0.897, *P* = 0.005) and appeared bimodal. Seventeen individuals had a mean fertility of 55% to <90% and 15 individuals had >90% to 100% fertility, respectively. Hence, the observed ratio of ‘low’ to ‘high’ fertility F_2_ plants (8.5 : 7.5) was not significantly different from a digenic epistatic ratio of 9 : 7 (χ^2^ = 0.127, d.f. = 1, *P* = 0.722). If the two fitted distributions were taken into account (see long-dashed vs. solid lines in Figure 6(B)), one individual from the ‘high’ class (marked by an asterisk) appeared to fall into the ‘low’ class, resulting in an exact ratio of 9 : 7 (N = 18 : 14; χ^2^ = 0.0, d.f. = 1, *P* = 1.0). Yet regardless of such ambiguity, these results are consistent with the hypothesis that reduced F_2_ pollen fertility (viz. ‘hybrid incompatibility’) between *S. englerianus* and *S. flavus* results from the interaction of two independently assorting genes. According to this ‘duplicate recessive- epistatic gene model’, one dominant allele *at each* gene would be sufficient to reduce fertility, while homozygosity for a recessive allele at *either* gene, or at *both* genes, causes high pollen fertility (Griffiths et al. 2000).

**Figure 6.**
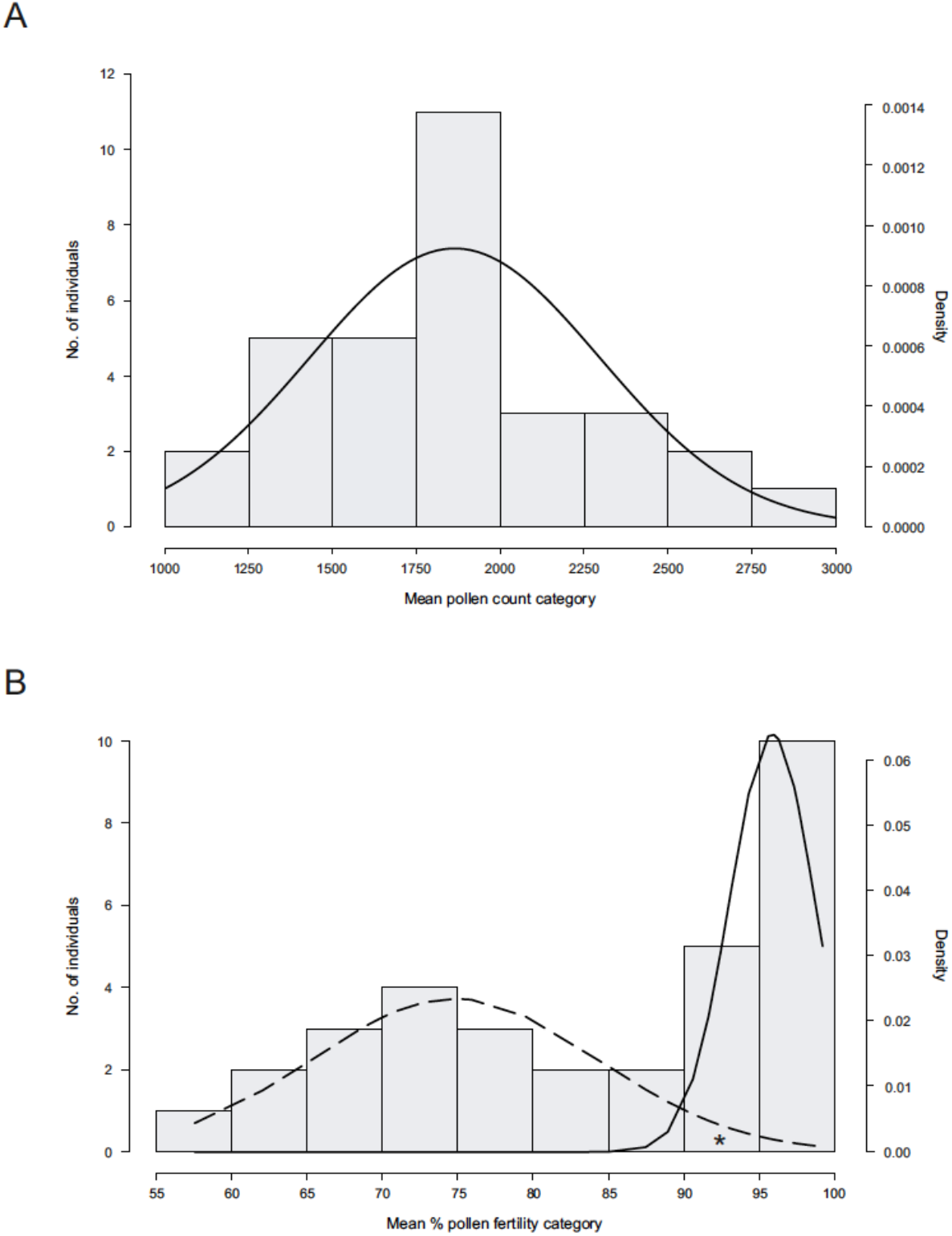
(A) Histogram of mean pollen counts for *Senecio englerianus* x *S. flavus* F_2_ hybrids (N = 32) with fitted normal distribution (solid line), using R. (B) Histogram of mean percentage pollen fertility in the same F_2_ generation. Smooth long-dashed and solid lines indicate expected F_2_ frequencies according to two normal distributions for the ’low’ and ’high’ F_2_ series, respectively, as fitted by the R package MIXTOOLS. Note, after curve fitting, one individual from the ‘high’ class (marked by an asterisk) appeared to fall into the ‘low’ class, resulting in an exact ratio of 9 : 7 (see text).

### Genetic basis of pappus type

The *S. flavus* individual (AS.1) used as the parent of the F_1_ and F_2_ generations produced achenes with connate-fluked (CF) pappus hairs (in addition to ‘typical’, non-fluked ones), while the *S. englerianus* parent (3.29) did not. Their F_1_ hybrid also produced achenes with a CF pappus. Of 87 F_2_ individuals, 68 exhibited this pappus type as well, while the remaining 19 produced only a non-fluked (NF) pappus. Hence, the CF : NF ratio observed (3.58 : 1) was not significantly different from a Mendelian 3 : 1 ratio (χ^2^ = 0.437, d.f. = 1, *P* = 0.509). These results are consistent with a single (‘major’) gene model, involving a completely dominant allele for expression of the CF pappus present in *S. flavus* and a recessive allele for the NF pappus at the locus in *S. englerianus*.

### Present and past climatic distribution predictions

The mean AUC training/testing values for the current distribution models of *S. englerianus* (0.999/0.999) and Southwest African *S. flavus* (0.994/0.994) were very high, indicating excellent predictive model performance (see Pflugbeil et al. 2021, and references therein). For *S. englerianus*, the ENMs for the present, the HCO (c. 6 kyr ago) and the LGM (c. 22 kyr ago; see Figures 7(A–C)) indicated relatively stable environments over time, with generally higher suitability (threshold c. ≥ 0.7) in the Central than Northern Namib Desert, especially during the LGM. Considering *S. flavus*, present and HCO models (Figures 7(D) and (E)) indicated most favourable conditions along the Namib Escarpment of western/southern Namibia (the species’ actual core-range) but also predicted areas of lower suitalbility (c. 0.3–0.7) where the species is either extremely rare (central/eastern Namibia, interior of South Africa) or absent (Northern Namib Desert). During the LGM, suitable habitat for *S. flavus* would have existed in the Central Kalahari Desert (Botswana), the Lake Victoria Region (LVR), the Horn of Africa, the Sahara Mts., and Northwest Africa (Figures 7(F) and (G)). This LGM model, however, failed to recover a continuous African Dry-Corridor (ADC) from Southwest Africa to Northeast Africa (see white-dashed lines in Figure 7(G)). The same was true when all 150 records of *S. flavus* from across the species’ range were taken into account (see Figure S3).

**Figure 7.**
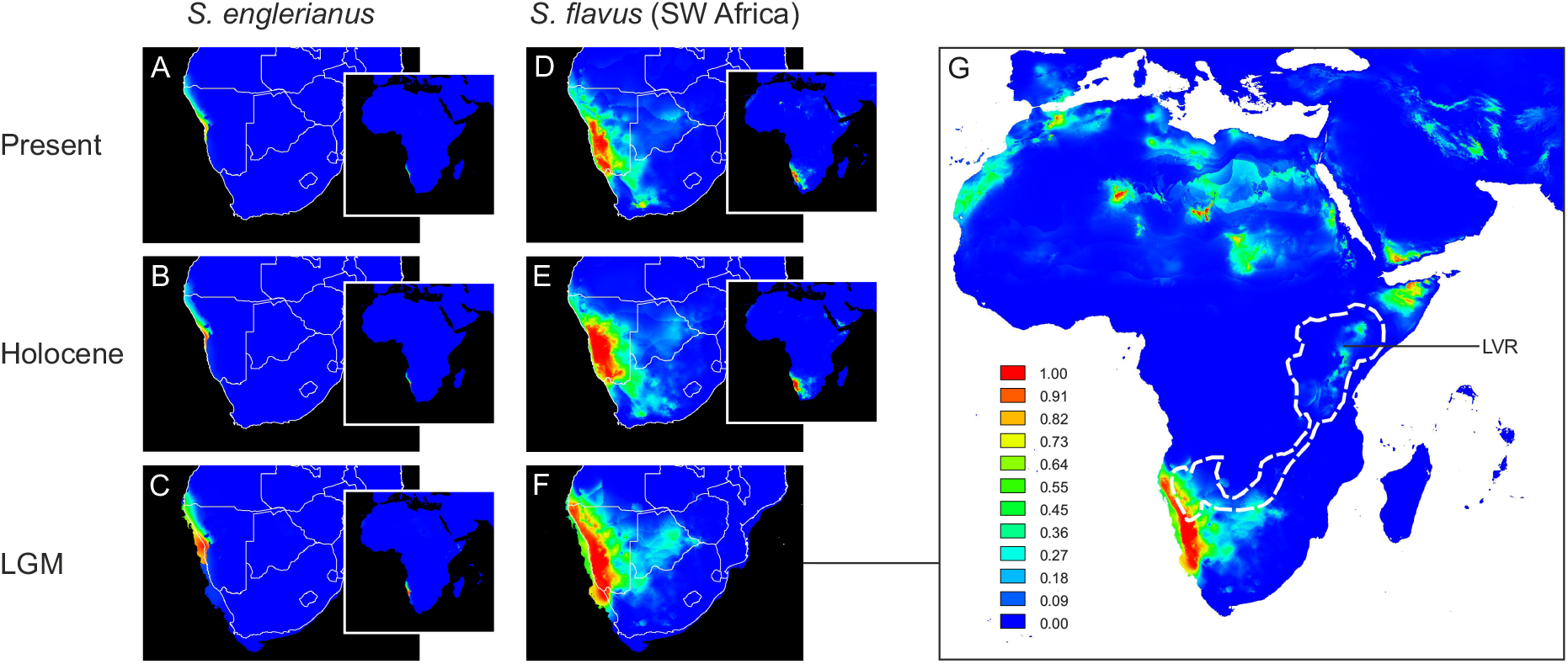
Ecological niche models (ENMs) for *Senecio englerianus* (A–C) and Southwest African *S. flavus* (D–G) in Namibia and adjacent regions at the present (1950*–*2000), the mid-Holocene Climate Optimum (HCO, c. 6 kyr ago), and the Last Glacial Maximum (LGM, c. 22 kyr ago). For each time period, potential distributions are also shown at the continental scale (see inserts A–E, and enlarged figure G for *S. flavus* at the LGM). ENMs were generated in MAXENT on the basis of 11 current bioclimatic variables and a total of 66 specimen records (*S. englerianus*: N = 30; *S. flavus*: N = 33; see text for details). Predicted distribution probabilities (ranging from zero to one) are shown in each 2.5 arc-minute pixel. In G, the white-dashed lines delimit the African Dry-Corridor (ADC) during arid periods of the Pleistocene according to Neumann and Scott (2018). LVR, Lake Victoria Region. See Figure S3 for corresponding ENMs of *S. flavus* based on all 150 locality records from across the species’ range.

## Discussion

### Evolutionary history of the ‘petiolate’ clade of S. englerianus and S. flavus

Our phylogenetic analyses of Senecioneae based on ITS data (Figures 3 and S1) clearly support the monophyly of the ‘petiolate’ clade of *Senecio englerianus* and *S. flavus*. They also confirm earlier phylogenetic estimates (Pelser et al. 2007) that the two target species are not only distantly related to sect. *Senecio* viz. the ‘core clade’ of *Senecio* (*sensu* Kandziora et al. 2017) but are neither part of *Senecio* s.str. Furthermore, age estimates indicate the stem age of the ‘petiolate’ clade possibly dates back to the early Late Miocene, c. 10.91 Mya (95% HPD: 23.14–3.26 Mya), whereas its crown age falls into the Early Pleistocene, c. 1.47 (2.72–0.11) Mya (Figure 3, Table S5). Clearly, these time estimates (especially for the stem age) have broad HPD intervals, and must be viewed cautiously. Also, despite the fact our taxon sampling covers c. 71% of all species of *Senecio* s.str. (15/21) reported for the Nambian Flora (Klaasen and Kwembeya 2013; Craven and Kolberg 2021), and c. 149 sub-Saharan (incl. 119 South African) species of Senecioneae (see Appendix 2 – *Table A2.1*), possibly more than 120 nominal species of *Senecio* are still missing from the ‘Afrotropic’ ecozone (Kandziora 2015; Kandziora et al. 2017). Hence, it remains unclear whether the long branch subtending the ‘petiolate’ clade reflects (1) an artifact of incomplete taxon sampling; (2) an extra-Namibian origin of this clade; or (3) long-term stasis and/or extinction of putative earlier lineages of *S. englerianus*/*S. flavus in situ*.

Nevertheless, our estimated stem age for this ‘petiolate’ clade remarkably coincides with the onset of cold north-bound currents along the southwestern coast of Africa (Benguela-Namib Upwelling, BNU) during the early Late Miocene (c. 12–10 Mya; due to an increase in strength of the Antarctic Circumpolar Current, ACC; c. 14 Mya), and which triggered (or intensified) arid or semi-arid conditions in today’s Namib Desert region (e.g. Heinrich et al. 2011; Rommerskirchen et al. 2011; Nylinder et al. 2016; Wan et al. 2021). By contrast, the clade’s much younger crown age slighty post-dates the strengthening and further (abrupt) cooling of the BNU system during the Early Pleistocene (c. 2.1 to 1.9 Mya; Marlow et al. 2000; see also Berger et al. 2002; Rommerskirchen et al. 2011). Molecular phylogenetic studies of other Namibian clades, though few in number, likewise report long ‘waiting times’ between origin and diversification, in both plants (e.g. *Indigofera*: Schrire et al. 2009; *Monsonia*: García-Aloy et al. 2017) and animals (e.g. insects, reptiles: Lamb and Bond 2013; Collins et al. 2019; Childers et al. 2021). In general, this decoupling of species richness from stem age is difficult to explain, but could reflect ecological limits to species richness (Stadler et al. 2014), combined with low (or heterogenous) rates of net diversification rather than high rates of extinction (Donoghue and Sanderson 2015).

Any intensification of BNU upwelling (i.e. increasingly cooler surface waters) during the Pleistocene would have increased xeric conditions in Southwest Africa, which in turn, likely forced local speciation events, for instance through the formation of aeolian sand dune systems, erosion along drainage lines, or climate-induced vegetation change (e.g. Marlow et al. 2000; Hoetzel et al. 2015). It is feasible, therefore, that the ‘petiolate’ clade of *S. englerianus*/*S. flavus* represents another example of ancient Namibian ‘relict’ clades that underwent relatively recent speciation in response to palaeoclimatic events over Pleistocene time scales. However, processes linked to diversification (speciation and extinction) in the Nambian Flora remain poorly understood and deserve further investigation (Linder 2014; García-Aloy et al. 2017).

### *Unresolved species relationships between* S. englerianus *and* S. flavus

Progenitor-derivative (P–D) species pairs are recognised as appropriate systems for studying plant speciation, as they represent instances of recent divergence, making it more feasible to identify presumably few inter-specific differences (e.g. in geography, ecology, morphology) and to infer possible early-evolving RI mechanisms (Crawford 2010). Such P–D pairs are often expected to display paraphyletic relationships, with accessions of the derivative species occupying a nested position within those of the ancestral taxon (e.g. Lopéz et al. 2012; Otero et al. 2019). However, our phylogenetic (ML) analysis of multiple intraspecific ITS sequences of *S. englerianus* and *S. flavus* (Figure S2) rsulted in an unresolved (polytomous) crown topolgy, including a weakly supported, potentially derived cluster of northern *S. flavus*. However, for recently diverged species, such failure to recover paraphyletic (P–D) relationships is not entirely unexpected for a number of reasons, including insufficent time for lineage sorting and hybridization/introgression (e.g. Mutanen et al. 2016).

Our experimental crossings demonstrated that *S. englerianus* and *S. flavus* are interfertile, with one F_1_ hybrid producing vigorous F_2_ offspring after self-pollination (albeit with variable pollen numbers and fertilities; Figure 6). This suggests that intrinsic postzygotic barriers to hybridization are relatively weak. However, there are no reports of naturally occurring hybrids between the two species in Namibia, suggesting that interbreeding there is largely prevented by prezygotic and extrinsic postzygotic barriers (see below). Hence, the species’ unresolved relationship (in terms of ITS) is more likely to reflect lack of sequence variation and/or insufficient time to lineage sorting rather than hybridization/introgression.

### *Evidence for species status of* S. englerianus *and* S. flavus

Despite a lack of separation in phylogenetic analyses, our study provides strong evidence that *S. englerianus* and *S. flavus* are ‘fully-fledged’ species that markedly differ in life history (short-lived perennial vs. annual), quantititative traits (Figure 5), breeding system (outcrossing vs. selfing), and pappus type (non-fluked vs. connate-fluked; Figures 2(E) and (F)). Notably, the two species occupy distinct ecoregions in Namibia (*S. englerianus*: Central/Northern Namib Desert; *S. flavus*: Nama-Karoo, Succulent Karoo and, rarely, Savanna), with only slight range overlap in the Central Namib Desert and its hinterland (approximately south of Mt. Brandberg to Khan River; see black rectangle in Figure 1(B)). According to our HUMBOLDT- derived PCA (Figure 4), *S. flavus* occupies higher-elevation sites with stronger fluctuations in annual temperature as well as higher (and more predictable) rainfall than *S. englerianus*. In fact, the corresponding niche overlap and divergence tests (NOT, NDT) revealed (near-) zero *D* values, suggesting that ecogeographical isolation (*sensu* Sobel et al. 2010) as well as adaptive niche divergence (Brown and Carnaval 2019) had an important role in speciation. The annual life form and less succulent leaves of *S. flavus* (Table 4) could thus reflect an adaptation to slightly more mesic conditions, as encountered in the Nama-Karoo relative to the hyper-arid Namib Desert.

In addition, there are three lines of evidence concerning mating type differences between the two species: (1) during cultivation, *S. englerianus* failed to set seed when left to self (JJ Milton, unpublished data), whereas *S. flavus* produces seed readily on selfing (see also Coleman et al. 2003); (2) in line with ‘selfing-syndrome’ theory (Ornduff 1969; Sicard and Lenhard 2011), the non-radiate capitula of *S. flavus* are significantly smaller than those of *S. englerianus* (Table 4), whereby the latter may occasionally produce ray florets (Jürgens et al. 2021) that could further help attracting pollinators; and (3) the *S. englerianus* individual examined for number of pollen grains per floret produced more than six times the amount of pollen seen in *S. flavus* (Table 3); this concurs with the notion that outcrossing species normally produce much more pollen per ovule than do selfers (Cruden 1977). Together, these results strongly suggest that *S. englerianus* is a self-incompatible, obligate outcrosser, whereas *S. flavus* is a self-compatible, predominantly autogamous species.

### *Evidence for* S. flavus *as a putative derivative of* S. englerianus *and the build-up of RI barriers*

A break-down of self-incompatibility, and/or shifts from outcrossing to selfing, have long been considered hallmarks of directional phenotypic evolution, facilitating the identification of putative derivative species and cases of ‘budding’ speciation (Crawford 2010; Anacker and Strauss 2014). In fact, unidirectional transitions from outcrossing to selfing are commonly reported in angiosperms (Takebayashi and Morell 2001; Igic and Busch 2013; Barrett 2014; Gamisch et al. 2015; although see Ferrer et al. 2006; Kim et al. 2008). Accordingly, it seems plausible that *S. flavus* was derived from *S. englerianus*. If so, the former could represent a rare example of a *wider-*ranged derivative that likely originated via parapatric (or ‘budding’) speciation at the periphery of a *smaller-*ranged progenitor (see also Kadereit 1984a; Kadereit et al. 1995). In support of this non-allopatric scenario, our palaeodistribution models suggest that the two species also occupied adjacent ranges in Namibia during the HCO (c. 6 kyr ago) and the LGM (c. 22 kyr ago), at which times (or earlier pluvial/arid periods) they might have even been broadly sympatric, especially in the Northern Namib Desert (see Figure 7). Although these models should be treated with caution (e.g. Wielstra 2019), they indicate that the species’ presently abutting ranges do not result from allopatric speciation followed by range expansion.

The ability of selfing variants to reproduce when pollinators or mates are rare or absent (Baker 1955) is generally favoured in small isolated or peripheral populations (Busch 2005; Iritani and Cheptou 2017). It is feasible, therefore, that the transition to selfing was among the early-acting reproductive isolation (RI) barriers involved in the origin of *S. flavus*. This prezygotic RI might have also accelerated the development of intrinsic postzygotic isolation (Fishman and Stratton 2004; see below). In addition, selfing would have increased the colonization ability of *S. flavus* (Baker 1955; Busch 2005), and possibly allowed this species to adapt more readily to its ‘novel’ (more mesic) environments (Coyne and Orr 2004), i.e. by blocking gene flow from *S. englerianus* at the time of speciation (see also Sobel et al. 2010). Morevover, in concert with the spread of selfing, other prezygotic (e.g. phenological) barriers could have played a role in the initial speciation process. However, there is no evidence from the herbarium material examined (collection Bertil Nordenstam) that the two species are isolated through differences in flowering time (both approximately January–July).

Local adaptation leading to ecogeographic isolation and niche divergence (see above) would be expected to impose a strong extrinsic postzygotic isolating barrier (Richards et al. 2016); hence, evidence from a crossing experiment should indicate some form of intrinsic postzygotic isolation between the two species (Coyne and Orr 2004; Sobel et al. 2010). Although our synthetic F_1_ and F_2_ material showed no signs of reduced vitality, the F_2_ exhibited a bimodal distribution of pollen fertilities (Figure 6(B)) compatible with a digenic-epistatic gene model, as expected for Dobzhansky-Muller (DM) incompatibilities (Fishman and Stratton 2004; Johnson 2008; Satogankas et al. 2020). In the context of parapatric speciation, such intrinsically lowered hybrid fitness is generally thought to arise as a ‘by-product’, i.e. after ecogeographical forms of isolation have evolved “to an appreciable level” (c.f. Sobel et al. 2010). In our system, therefore, this type of ‘hybrid incompatibility’ is unlikely to have contributed to speciation, and might (at present) contribute to RI only in the species’ immediate contact zone in the Central Namib Desert region (see black rectangle Figure 1(B)). Nonetheless, our inference of DM incompatibilities could point at marked adaptive-genomic divergence between *S. englerianus* and *S. flavus*, related to their pronounced niche divergence in Namibia.

### Joint evolution of key traits in *S. flavus* affecting adaptation capacity and dispersal

While selfing would have allowed *S. flavus* to successfully colonise unpredictable and ‘novel’ habitats, its heritable ability or *propensity* to disperse (*sensu* Billiard and Lenormand 2005) was likely facilitated by the evolution of connate-fluked (CF) pappus hairs (Figure 2(F)). It has been long suggested that the grapple-like tips (or ‘prongs’) of these hairs, which remain firmly attached to the achene (RJ Abbott, pers. obs.), facililate adhesive animal dispersal (Drury and Watson 1966; Jeffrey 1979; Lawrence 1981; Coleman et al. 2003), hence in addition to dispersal by wind afforded by the non-fluked (‘typical’) hairs that are exclusively present in *S. englerianus* (Figure 2(E); Coleman et al. 2003). Notably, of the 245 species of *Senecio* examined by Drury and Watson (1966), c. 40 possessed ‘fluked’ pappus hairs, i.e. with inverted or retrorse teeth. However, fluked pappus hairs with conspicuous ‘prongs’, as observed in *S. flavus*, seem to be rare in the genus (e.g. *S. californicus* DC.; Drury and Watson 1966). Our genetic analysis demonstrated that this complex hair structure is controlled by a single major gene with completely dominant allele expression, adding weight to the view of Coleman et al. (2003) that it arose as a new mutation in *S. flavus*. Mutations of such large-effect loci are generally expected to be favoured during adaptive species divergence with gene flow (e.g. Tigano and Friesen 2016). Accordingly, it is tempting to speculate that *S. flavus* represents an example of ‘budding’ speciation where selection favoured a combination (or even tight linkage) of major genes controlling not only dispersal propensity (CF pappus) but also reproductive ability (selfing) and a fast life cycle (annuality), i.e. three key traits jointly affecting adaptation capacity and invasiveness (e.g. Kandziora et al. 2017; Beckman et al. 2018).

### *Northward range expansion of* S. flavus

Coleman et al. (2003) proposed that the CF pappus had a pivotal role in the long-distance dispersal (LDD) of *S. flavus* from Southwest to North Africa by facilitating the attachment of achenes to ground-feeding birds migrating annually between these regions (e.g. *Motacilla* spp., *Lanius* spp.). Based on ITS data, these authors inferred a Late Pleistocene origin for the Southwest–North African split of *S. flavus* (c. 0.15 Mya; i.e. based on the single informative sequence position also inferred herein). In contrast, using isozyme data, Liston et al. (1989) dated this split back to the Early Pleistocene (c. 0.77 Mya) and attributed this disjunction to vicariance, i.e. range contraction following the species’ northward expansion along the African Dry-Corridor (ADC). Consistent with either time estimate above, this potential migration route is generally believed to have reached its greatest development during arid (i.e. high-latitude glacial) periods of the Pleistocene. At these times, this ‘arid track’ likely extended in a northeasterly direction south of the Congolian Rainforest Basin into the Lake Victoria Region (LVR) of East Africa (see white-dashed lines in Figure 7(G)), repeatedly joining the xeric grassland and savanna vegetation of the Namib–Karoo–Kalahari region with the Saharan via the Horn of Africa (van Zinderen Bakker 1975; Neumann and Scott 2018).

In fact, the ADC is frequently invoked in the floristic and biogeographic literature (e.g. Winterbottom 1967; Verdcourt 1969; de Winter 1966, 1971; White 1983; Jürgens 1997; Banfield et al. 2011; Linder 2014; Perkins 2020), despite controversy over its dates of establishment and closure (e.g. Balinsky 1962; Thiv et al. 2011; Bellstedt et al. 2012; Pokorny et al. 2015; Kissling et al. 2016; Kandziora et al. 2017). However, to our knowledge, no previous study has explicitly examined (e.g. modelled) the temporal and spatial dynamics of this corridor for a particular plant (or animal) species. Our distribution model of Southwest African *S. flavus* for the LGM (Figures 7(F) and (G)) reconstructed part of the predicted ADC at its northeastern end (in the LVR) with c. 0.3 distribution probability, and more suitable habitats (c. 0.5–1.0) throughout the Horn of Africa, across isolated Saharan mountain ranges (e.g. Ennedi Plateau, Tibesti/Hoggar Mts.), and in Northwest Africa (West Sahara, Rif Mts.). Our results further show that the climate conditions of the LGM (and likely earlier arid periods) could have favoured the northeastward expansion of *S. flavus* from Namibia into the Central Kalahari Desert (Botswana); however, this does not seem to have resulted in a contiguous corridor (Figure 7(G)), likely because conditions were still too humid for this (semi-)desert annual in areas south of the Congolian Basin (Collins et al. 2011, 2014; Neumann and Scott 2018). Our ENM results, therefore, do not exactly match the expected ADC pattern, but nonetheless raise the possibility that the Southwest–North African disjunction of *S. flavus* was established by dispersal across intermediate areas rather than intra-continental LDD.

### Evolutionary ramifications associated with the northward spread of *S. flavus*

*Senecio flavus* is known to have given rise to two polyploid taxa in North Africa through hybridization with an annual diploid member of the ‘*vulgaris*-clade’ of *Senecio* s.str., namely *S. glaucus* L. subsp. *coronopifolius* (Maire) Alexander (Kadereit et al. 2006, and references therein). These are the hexaploid *S. hoggariensis* Batt. & Trab. (2*n* = 6x = 60) and the tetraploid *S. mohavensis* A.Gray ssp. *brevifolius* (Kadereit) M.Coleman (2*n* = 4x = 40), originally described as a subspecies of *S. flavus* (Kadereit 1984b) and now known to be distributed from the Sinai Peninsula eastwards to Pakistan (Coleman et al. 2001). In turn, *S. mohavensis* ssp. *breviflorus* has been implicated as progenitor of ssp. *mohavensis* (2*n* = 4x = 40), which is endemic to the Mojave and Sonoran Deserts of North America (Liston et al. 1989; Coleman et al. 2001, 2003).

Hence, there is now mounting evidence to support a step-like biogeographic scenario (Figure 8) where (1) *S. flavus* originated in the Namib Desert region; (2) then spread to North Africa, where allopolyploidy (involving *S. glaucus*) not only increased species diversity (*S. hoggariensis*), but could also have triggered or facilitated eastward range expansion (*S. mohavensis* ssp. *brevifolius*; Comes and Abbott 2001; Kadereit et al. 2006); and (3) cross-continental LDD and intraspecific differentiation in southwestern North America (ssp. *mohavensis*; Coleman et al. 2003). When combined with our estimated crown age of the ‘petiolate’ clade (c. 1.47 Mya), this scenario testifies to the long-held notion (e.g. Stebbins 1952; Axelrod 1972) that plant speciation may proceed rapidly in extreme (i.e. hyper-arid to semi-arid) environments over Pleistocene time scales (see also Kadereit and Abbott 2021).

**Figure 8.**
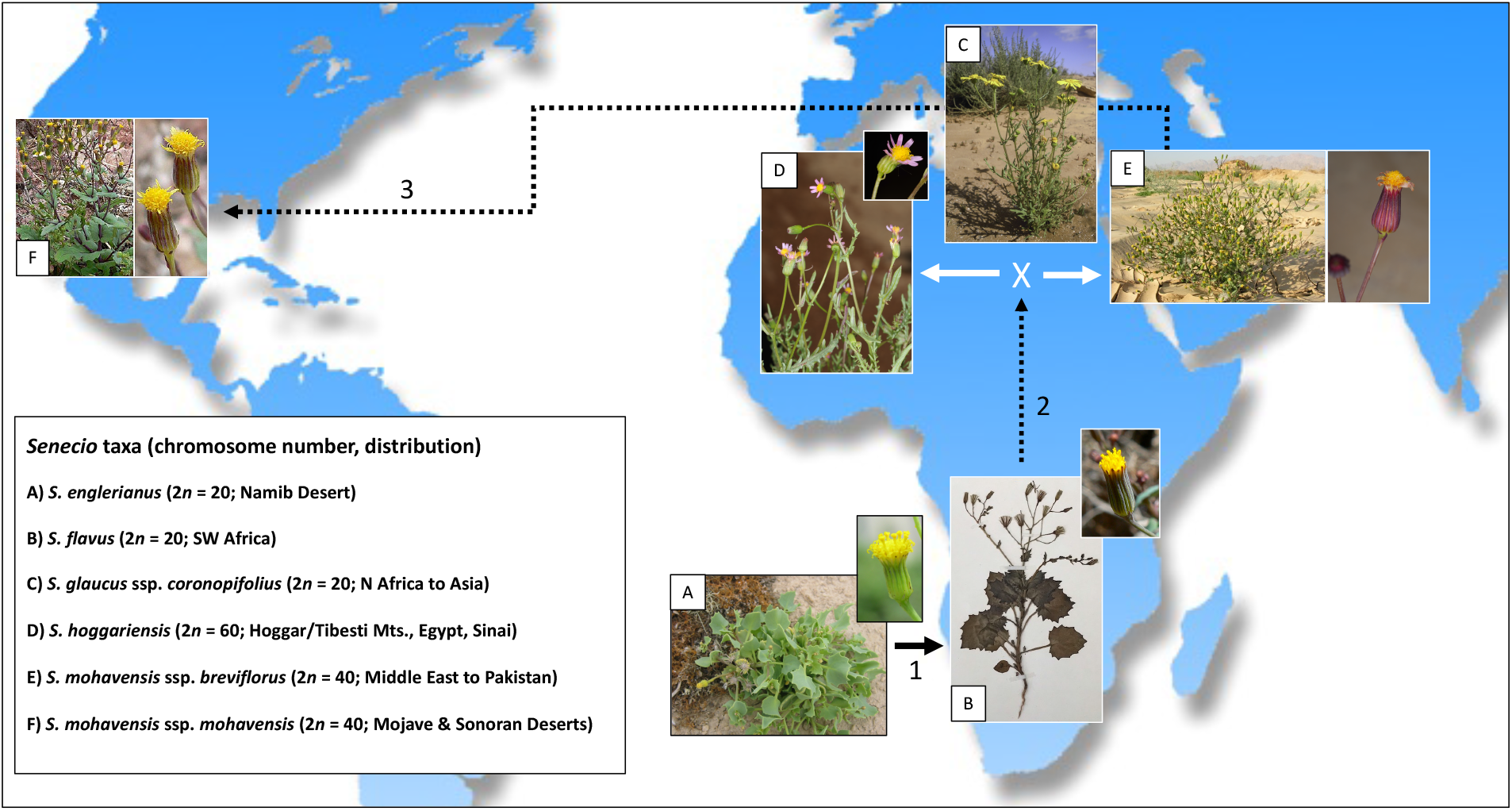
Hypothesized, step-like biogeographic scenario involving (1) the origin of *Senecio flavus* from *S. englerianus* in the Namib Desert region, followed by (2) its spread to North Africa (Liston et al. 1989; Coleman et al. 2003; this study), where hybridization with *S. glaucus* ssp. *coronopifolius* gave rise to *S. hoggariensis* and *S. mohavensis* ssp. *breviflorus*, respectively (Kadereit et al. 2006). Note the cross-continental dispersal (3) of the latter taxon to southwestern North America (Coleman et al. 2003), followed by the origin of ssp. *mohavensis*. Photographs by Joseph J. Milton (A: habitus), David Forbes (A: close-up); Richard J. Abbott (B: habitus, collection Bertil Nordenstam), Fouad Msanda (B: close-up; CC BY-NC 4.0), Philippe Geniez (C; BY-NC 4.0), Ori Fragman-Sapir (D; and E: close-up; www.flora.org.il); Mori Chen (E: habitus; www.flora.org.il), and Stan Shebs (F; CC BY-SA 4.0). Worldmap (Wagner II projection) by Tobias Jung (CC BY-NC 4.0).

Perhaps even more remarkably, the reticulate evolution of *S. flavus* with *S. glaucus* in North Africa involved the exchange of (both plastid and nuclear) genomes between species that, according to our ITS chronogram (Figure 3), last shared a common ancestor c. 12.7 million years ago. This inference is intriguing since hybridisation in plants (and animals) is generally thought to predominantly occur between recently diverged species (e.g. Mallet 2005; Paun et al. 2009; Abbott 2013, 2017). On the other hand, there are occasional examples of long-diverged yet still hybridising species, and likely more so in plants than in animals (e.g. Rieseberg 2000; Edmands 2002; Coyne and Orr 2004; Rothfels et al. 2015; Ebersbach et al. 2020). However, why genetic (or hybrid) incompatibilities evolve at different rates, and not always accumulate linearly (or even faster) with time as parental lineages diverge, remains poorly understood (e.g. Orr and Turelli 2001; Paun et al. 2009; Wang RJ et al. 2013, 2015; Kulmuni et al. 2020).

### Conclusions and future directions

Our results provide support for a rare example of parapatric (‘budding’) speciation in which a *wider-*ranged derivative (*S. flavus*) originated at the periphery of a *smaller*-ranged progenitor (*S. englerianus*) in the Namib Desert region of Southwest Africa. Together with ecogeographical isolation and adaptive niche divergence, shifts to selfing and annuality as well as higher dispersal propensity may all have contributed to this speciation event, and ultimately the ability of *S. flavus* to expand to the (semi-)desert areas of North Africa, perhaps by using parts of the African Dry-Corridor as ‘stepping stones’ to reach East Africa during arid periods of the (Late) Pleistocene. That said, further phylogenetic sampling is now required to evaluate the hypothesized ‘relict status’ of this ‘petiolate’ clade, and whether it represents a relictual and likely separate genus. In addition, population-level genomic analysis remains to be done to precisely date the split between *S. englerianus* and *S. flavus* (and between southern and northern populations of the latter species), ideally in combination with high-resolution palaeodistribution modelling over extended (i.e. full Plio-/Pleistocene) time scales (Gamisch 2019).

Finally, more detailed information (e.g. genomic, ecological, morphometric, experimental-reproductive) is desirable for this species pair in Southwest Africa. In particular, whole genome sequence comparisons should shed light on the genetic basis of adaptive divergence between the two species and how this might associate with the emergence of intrinsic postzygotic reproductive isolation between them (e.g. Hu et al. 2021; Wan et al. 2021; Wilkinson et al. 2021). Taken overall, we hope this paper helps motivate the development of purposefully integrated research programs to further study processes of speciation not only in this specific *Senecio* system, but also in other closely related (ideally sister) species of the Namibian desert vs. inland flora (e.g. Maggs et al. 1998; Craven and Craven 2000). We believe a final answer to the biogeographic importance of the African Dry-Corridor will emerge only after analysis of many genera in Southwest Africa with close relatives in North and/or East Africa (e.g. *Camptoloma*, *Fagonia*, *Heliotropium, Kissenia*, *Zygophyllum*; Craven and Vorster, 2006; Craven 2009; Bellstedt et al. 2012; Pokorny et al. 2015).

## Acknowledgements

We are grateful to Chris Cupido (Fort Hare University) for help with field work in Southwest Africa, Bertil Nordenstam (Swedish Museum of Natural History) and Michael Möller (Royal Botanic Garden Edinburgh, RBGE) for advice and providing herbarium specimens, David Forbes and Ailan Wang (St Andrews University) as well as Frieda Christie (RBGE) for laboratory assistance, Pieter Pelser (University of Canterbury) for correspondence and preliminary trees of Senecioneae, Sarah Sophie Brandauer (Salzburg University) for help with mapping occurrence records, Max Coleman (RBGE) for providing SEM-pappus photographs (Figures 2(E) and (F)), Hans-Joachim Esser (Munich Herbarium) for having forwarded an important publication on the Namibian Flora, and Joachim Kadereit (Mainz University) for advice and very helpful comments on an earlier version of this manuscript.

## Data availability statement

Data on locality records, voucher specimens (Nordenstam collection) and ITS sequence alignments are avialibale upon request to

## Funding

This research was funded in part by the award of a Natural Environment Research Council (NERC) CASE research studentship to JJM.

## Supplementary material

**Figure S1.**
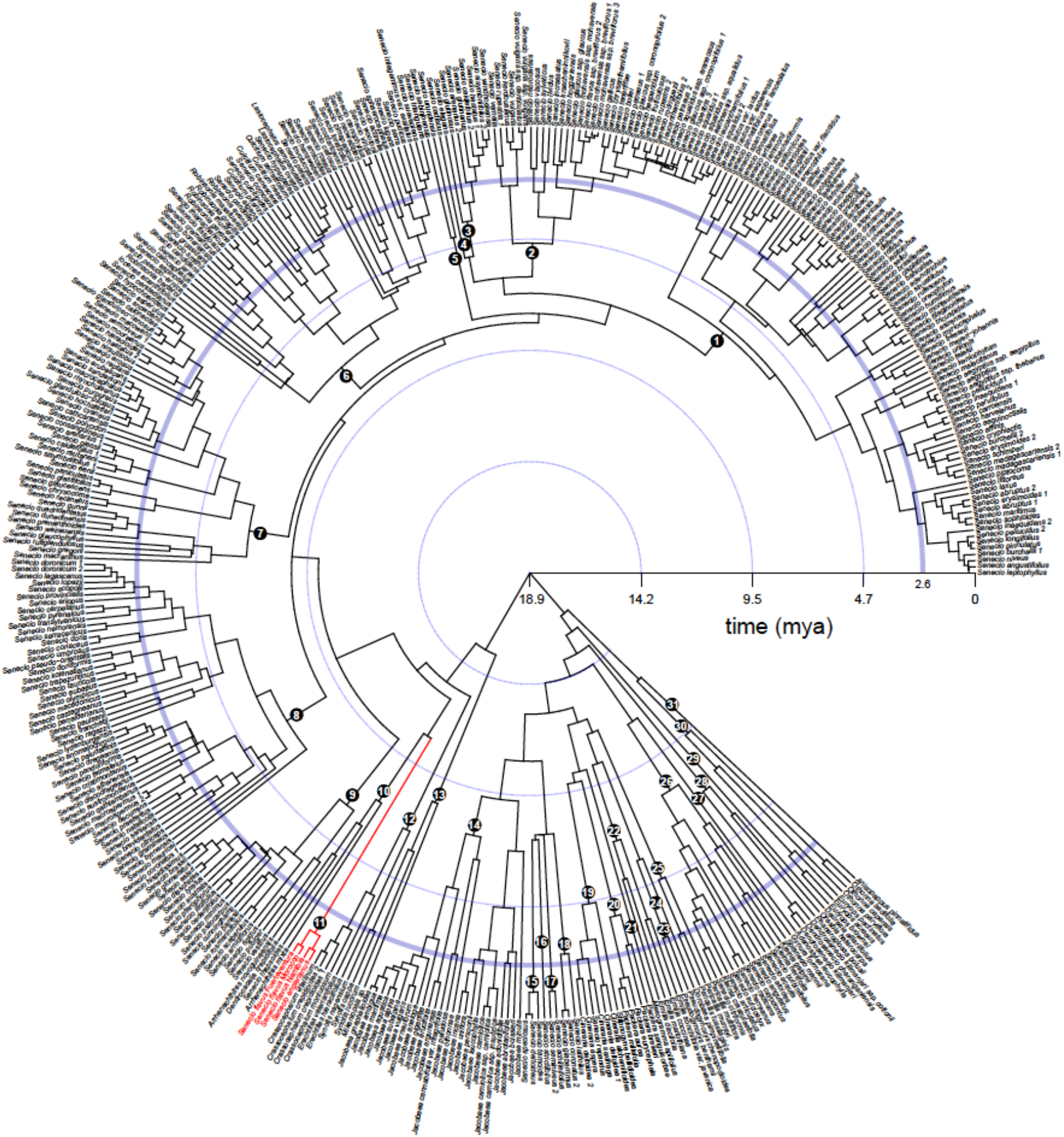
Full BEAST-derived maximum clade credibility (MCC) tree of Senecioneae, plus one member of Athroismeae (*Anisopappus pinnatifidus*), based on ITS sequence data (403 accessions, c. 363 spp.), and associated with Figure 3. Numbers of major clades and lineages are identified in Appendix 2 (Tables 2.1 and 2.2). The ‘petiolate’ clade of *Senecio englerianus*/*S. flavus* is highlighted in red. The blue, wider inner circle marks the beginning of the Pleistocene, c. 2.6 million years ago (Mya).

**All supplementary files are available on request**

**Appendix 1**: Methods. DNA extraction, ITS amplification, sequencing and cloning.

**Appendix 2:** *Tables A2.1–3*. Taxa included in the BEAST analysis of Senecioneae, using ITS sequence data (Figures 3 and S1).

**Table S1**: Description of bioclimatic variables retained for environmental niche analysis of *Senecio englerianus* and *S. flavus* in Southwest Africa (HUMBOLDT; Figure 4) and the species’ range-wide ecological niche modelling (MAXENT; Figures 7 and S3).

**Table S2**: Description of 14 trait variables measured on *Senecio englerianus, S. flavus* and F1 hybrids, together with loadings and variable contributions (%) for the first two principal components.

**Table S3**: Full matrix of quantitative trait data.

**Table S4**: Estimates of pollen number and fertility in *Senecio englerianus*, Moroccan *S. flavus*, one of their F_1_ hybrids, and the F_2_ generation.

**Table S5**: Stem and crown ages of major clades and lineages of Senecioneae based on the ITS chronogram (Figure 3).

**Figure S2**: Maximum likelihood (ML) phylogram for ITS sequences of the ‘petiolate’ clade, plus outgroup taxa.

**Figure S3**: Present and past ecological niche models (ENMs) of *Senecio flavus* based on all 150 locality records from across the species’ range.

## Notes

### Competing Interest Statement

The authors have declared no competing interest.

